# Inter- and intra-individual variability in structure-function coupling in human brain

**DOI:** 10.64898/2026.02.25.707886

**Authors:** Alina Studenova, Felix Ströckens, Luke J. Edwards, Anna-Lena Stroh, Saskia Helbling, Burkhard Maess, Kerrin J. Pine, Cam-CAN, Katrin Amunts, Evgeniya Kirilina, Nikolaus Weiskopf, Arno Villringer, Vadim Nikulin

## Abstract

Microstructure shapes functional brain dynamics. Yet this notion has remained largely untested for human brain signals measured with electroencephalography and magnetoencephalography (EEG/MEG). In this study, we related functional activity variance to microstructural variance across brain regions and across participants. Across regions, we used multiple independent datasets: EEG (n=209), MEG (n=507), high-resolution MRI (n=10), and the human post-mortem brain. Across participants, we used a multimodal dataset including MEG and high-resolution MRI (n=31). Between regions, alpha power correlated positively with layer IV thickness and with myelin and iron estimates at mid-cortical depth. By contrast, between participants, alpha power correlated negatively with mid-cortical myelin and iron estimates. Using computational modeling, we demonstrated that regional effects are consistent with excitatory changes, whereas inter-individual effects may be driven by inhibitory influence. This dissociation reveals distinct underlying mechanisms and indicates that different factors influence intra-versus inter-individual variations in alpha activity.

## Introduction

Macroscopic functional recordings, such as magnetoencephalography and electroencephalography (EEG/MEG), are used to establish the link between human brain dynamics and cognition in the living human brain (Klimesch, 1997; Mahjoory et al., 2019; Cesnaite et al., 2023). However, our understanding of the mechanisms underlying the generation of signals at the level of cells and cortical cellular architecture remains limited. The cortical cellular architecture (cytoarchitecture) has been investigated for a long time and with considerable success (Amunts et al., 2007; Shepherd et al., 2017), in both animal models and post-mortem human brain samples. To achieve a more comprehensive understanding of brain organization, the microscopic and macroscopic lines of research should be integrated, and anatomical and physiological properties need to be linked.

The basis of EEG/MEG signals is electrical and magnetic fields emerging from the synchronous activity of large populations of neurons (Schomer & Da Silva, 2012; Ilmoniemi & Sarvas, 2019). Excitatory pyramidal neurons in the neocortex are the main contributors to the observable EEG/MEG signal, due to the aligned orientation of their dendritic trees, forming directional currents and fields (Ilmoniemi & Sarvas, 2019). In contrast, inhibitory interneurons’ activity produces a closed field without directional coherence (Hagen et al., 2018), and thus is not observable at a distance. While there is tentative consensus that pyramidal neurons’ activity can be detected by EEG/MEG, understanding how pyramidal neuron density, laminar organization, and non-pyramidal neurons’ activity shape EEG/MEG signals is limited. Such an understanding is essential to accurately link these macroscopic signals to the underlying microcircuit dynamics and cortical cytoarchitecture to interpret interindividual differences within the healthy population and in patients with neurological conditions.

Cell distribution, i.e., cytoarchitecture, can be best measured in histological sections of postmortem brains. A notable milestone in post-mortem brain cytoarchitecture is BigBrain (Amunts et al., 2013). This ultrahigh-resolution 3D model of the human brain consists of 7404 histological sections stained for cell bodies at a resolution of 20 micrometers. The cytoarchitecture can, to a certain extent, be approximated from high-resolution magnetic resonance imaging (hMRI; also known as *in-vivo* histology MRI). Recent improvements in MRI image quality and biophysical modeling of the MRI signal provide a tentative link between structural macroscopic images and microstructure (Weiskopf et al., 2021). Multiparametric maps (MPMs) derived from MRI are sensitive to the myelin distribution and iron deposition (Edwards et al., 2018; Weiskopf et al., 2021), and myeloarchitecture is closely linked to cytoarchitecture (Dinse et al., 2015). Specifically, myelin content correlates with cell body size (Foit et al., 2022) and with cell count in layers II, III, IV, and VI (McColgan et al., 2021). This implies that depth-resolved myelin-sensitive parameters obtained non-invasively from living humans may serve as an approximation of cytoarchitecture at the resolution of individual laminae.

It has been shown that electrical activity in the neocortex is strongly influenced by its layered organization (for a comprehensive overview, see Table S1). In a computational model (Neymotin et al., 2020), the shape and frequency of the simulated alpha rhythm closely matched empirical data when pyramidal neurons were positioned in layers II, III, and V (layers that, in the real brain, contain the majority of pyramidal cells) and when the network received both feedforward and feedback inputs. In slices from rat visual cortex, alpha rhythm power was correlated with bursting in layers IV and V (Traub et al., 2020). In monkeys, local pacemakers of alpha rhythm were found both in superficial and deep layers of visual areas (Bollimunta et al., 2008; Haegens et al., 2015), as well as somatosensory and auditory areas (Haegens et al., 2015). However, communication between regions in the alpha range seems to occur in deep layers (Buffalo et al., 2011). In human patients with epilepsy, alpha rhythm was found to have high power primarily in supragranular layers (layer I, II, and III of the neocortex; Halgren et al., 2019). Apart from cortical generators, thalamic input plays a role in generating alpha oscillations, which mostly arrives at layer IV, or the granular layer (Shepherd et al., 2017). The alpha rhythm power over occipital sites was correlated with an increase in the signal from the thalamus, based on simultaneous fMRI-EEG (Goldman et al., 2002; Moosmann et al., 2003), while alpha power across the cortex was reduced when metabolism was reduced in the thalamus based on positron emission tomography (PET) (Lindgren et al., 1999; Schreckenberger et al., 2004). Furthermore, the power of alpha oscillations was found to increase with decreasing thalamocortical connectivity caused by white matter degeneration (Kumral et al., 2022).

Overall, research on the laminar distribution of brain dynamics can be broadly divided into three types. 1) **Inter-laminar studies** involve electrophysiological data primarily from animal models derived from small cortical patches of a cortex (similarly, detailed *in-silico* models typically also simulate a single cortical region). Although this group of studies has provided the most insights, recordings are often limited to the sensory cortices. Consequently, generalization to other cortical areas remains uncertain, given the substantial difference in structure and function between sensory and association regions. 2) **Inter-regional studies** use large datasets to correlate multiple features simultaneously (all functional and cytoarchitectonic characteristics; Shafiei et al., 2023). These studies employ a data-driven approach, but study correlations of averaged metrics from different modalities across brain regions. Often, they do not involve the same participants for electrophysiological and structural data, which considerably restricts the interpretation of the results. 3) **Inter-individual studies** combine MRI-EEG or fMRI-EEG studies in humans and use gross anatomical features. These studies take individual differences into account but lack the spatial resolution to determine the fine-grained laminar structure of the neocortex (Wagstyl et al., 2018).

In this study, we addressed these limitations by combining all three approaches in a single study, taking advantage of multimodal recordings to establish a link between brain microstructure and large-scale macroscopic brain dynamics (Fig. 1; the study was preregistered, Studenova, 2023). Our first approach was to explore across-region relationships between different characteristics of four recording modalities: EEG (LEMON, 209 participants; Babayan et al., 2019), MEG (Cam-CAN, 507 participants; Shafto et al., 2014; Taylor et al., 2017), 7T MRI (10 participants; McColgan et al., 2021), and high-resolution scans of stained histological slices from a postmortem brain (BigBrain, Amunts et al., 2013). While this addresses limits with respect to whole brain analysis and laminar specificity, it would still involve comparisons of different individuals across modalities. In order to overcome that limitation, our second approach was to focus on between-participant correlations using MEG and 7T MRI data, recorded from the same participants.

**Figure 1.**
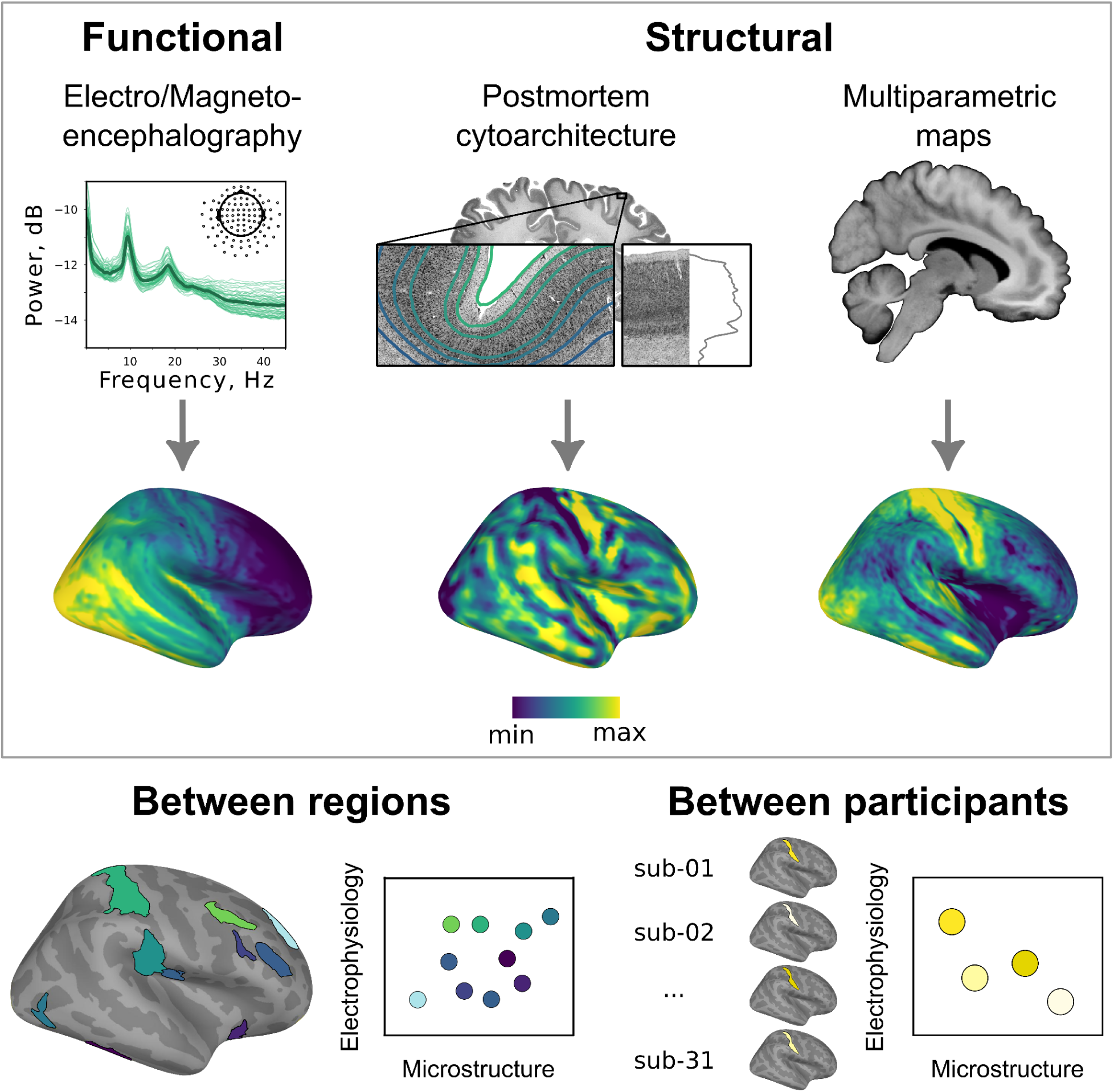
Methodological overview. We used different datasets of several modalities: the LEMON dataset with resting-state EEG (209 participants; Babayan et al., 2019), the Cam-CAN dataset with resting-state MEG (507 participants; Shafto et al., 2014; Taylor et al., 2017), BigBrain with cytoarchitecture (Amunts et al., 2013), the 7T MRI dataset with multiparametric maps (MPMs) (10 participants; McColgan et al., 2021), and our newly collected dataset with combined resting-state MEG and 7T MPMs (31 participants). From EEG/MEG, we derived the following variables (motivated by previous studies on laminar distribution of brain dynamics, see Table S1): theta power, alpha power, alpha frequency, low beta power, high beta power, 1/f slope estimated in the 2–40 Hz range, 1/f slope estimated in the 40–60 Hz range, nonsinusoidality of alpha rhythm amplitude (delta CT; Schaworonkow et al., 2019), and non-zero-mean of alpha amplitude (baseline-shift index; BSI; Nikulin et al., 2010). From the postmortem brain, we derived the following variables: thickness of supragranular, granular, and infragranular layers, and the mean and skewness of staining profiles. From structural 7T MPMs, we derived: myelin estimates in superficial, mid-cortical, and deep layers as measured by R1 and R2*. We looked at the associations in two ways: 1) between vertices (or regions), averaged over participants’ brains from different recording modalities, and 2) between participants within a vertex (or a region).

We found that correlations between regions (approach I) and between participants (approach II) led to opposite signs of correlation between microstructure and electrophysiology. Based on previous research, we hypothesized that this discrepancy may stem from the different contributions of excitatory and inhibitory neurons activity to inter-individual and between-region variability in brain structure. Biophysical modeling with two populations (excitatory and inhibitory) demonstrated the plausibility of this hypothesis.

## Results

### Selection of variables

For the between-region comparison, we used two different datasets of resting-state electrophysiological recordings: the LEMON dataset with EEG data (eyes open and eyes closed conditions; Babayan et al., 2019) and the Cam-CAN dataset with MEG data (eyes closed condition; Shafto et al., 2014; Taylor et al., 2017). Since we did not aim to study differences between eyes open/closed conditions, we integrated both to increase statistical power (the spatial distribution in our data was similar between the two conditions, see Fig. S1). We looked for spectral variables that exhibited maximal similarity in inter-regional distribution between MEG and EEG modalities (Table 1, step 1). We preregistered the correlation of 0.60 to be the lower threshold for the variable to be considered to have a similar whole-brain distribution in two modalities. The following variables were assessed: theta power, alpha power, alpha frequency, low beta power (15–20 Hz), high beta power (20–30 Hz), 1/f slope estimated in the 2–40 Hz range, 1/f slope estimated in the 40–60 Hz range, nonsinusoidality of alpha rhythm amplitude (delta CT; Schaworonkow et al., 2019), and non-zero-mean of alpha amplitude (baseline-shift index BSI; Nikulin et al., 2010). Whole-brain distributions of those variables for both datasets are shown in Fig. 2 and Fig. S1. The correlation of distributions exceeded the preregistered threshold (*r* = 0.60) for only three variables: alpha power, low beta power, and high beta power (correlations between EEG eyes closed and MEG eyes closed: theta power 0.278, alpha power 0.788, alpha frequency 0.591, low beta power 0.62, high beta power 0.667, 1/f slope 2–40 Hz 0.463, 1/f slope 40–60 Hz 0.598, nonsinusoidality of alpha rhythm amplitude 0.146, and baseline-shift index 0.187). Therefore, we conducted further analysis only on alpha power, low beta power, and high beta power.

**Figure 2.**
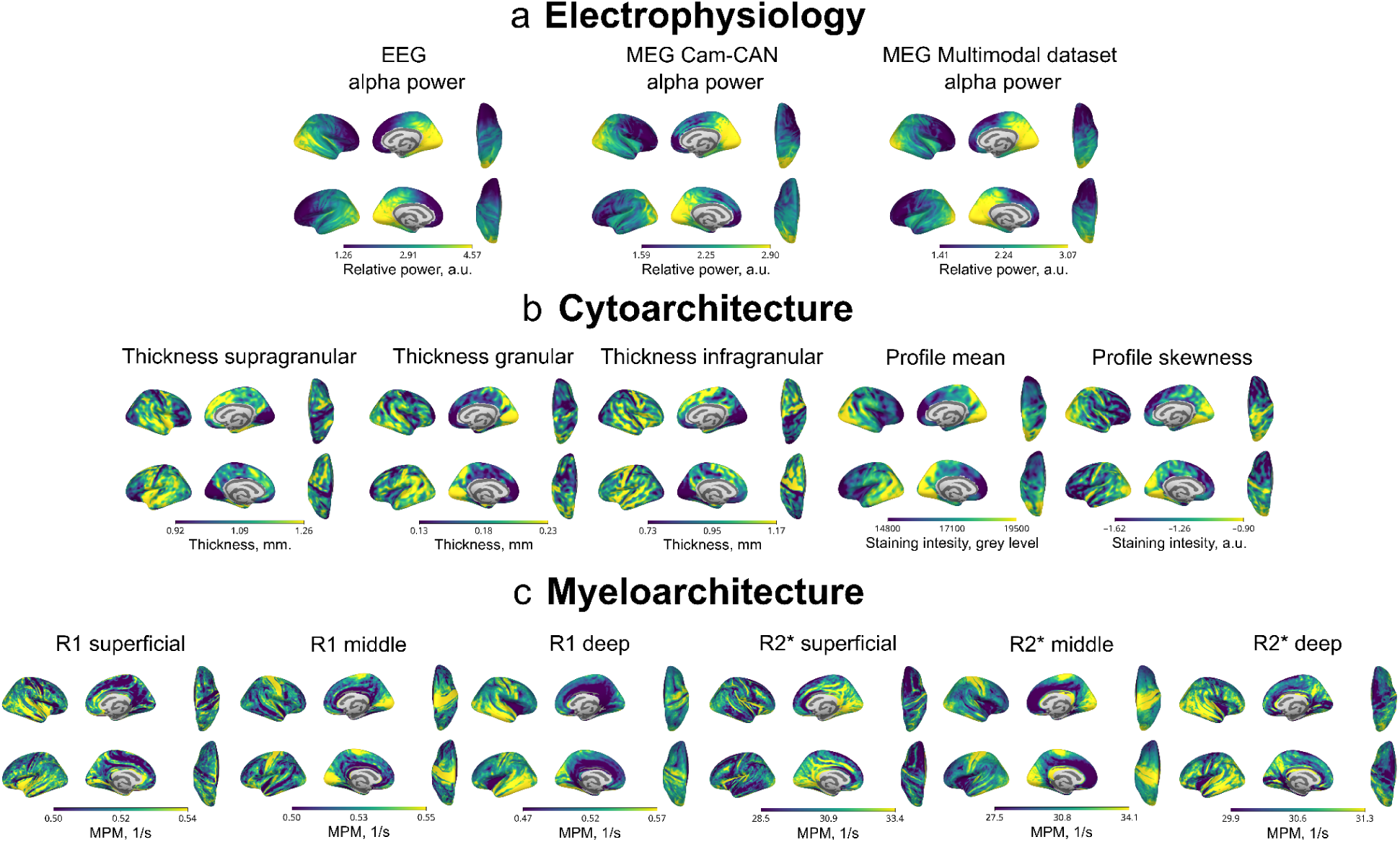
Distributions of the investigated variables (see also Fig. S1). **a**. Electrophysiological variables from EEG (LEMON) and MEG (Cam-CAN and Multimodal dataset) data. Alpha power was computed as the power ratio (relative power) P_pk_/P_ref_, where P_pk_ is the power ± 2 Hz around the peak frequency and P_ref_ is the average power in the two 1Hz intervals below and above the P_pk_ range. **b**. Cytoarchitectonic variables from BigBrain (post-mortem). We derived the thickness of each cortical layer, mean staining intensity across layers (which roughly correlates with the total amount of cells in a vertex), and skewness of staining intensity across layers (which correlates with the amount of cells and cell size in the supragranular layer in comparison with infragranular layers). Thicknesses were combined into supragranular (sum of layers I, II, and III) and infragranular (sum of layers V and VI). **c**. Multiparametric map parameters sensitive to myelin from 7T structural MRI (Multimodal dataset). Multiparametric maps (MPMs) were used to derive myelin estimates. We obtained approximations of myelin content as measured with R1 and R2* at three cortical depths (superficial, mid-cortical, and deep) using factor analysis (see Methods).

**Table 1.** Overview of preregistered analyses and outcomes (Studenova, 2023). Steps are described in succession as they were taken. Results from one step often, but not always, informed the analysis of the next step. We used the following datasets: the LEMON dataset with resting-state EEG (209 participants; Babayan et al., 2019), the Cam-CAN dataset with resting-state MEG (507 participants; Shafto et al., 2014; Taylor et al., 2017), BigBrain with cytoarchitecture (Amunts et al., 2013), the 7T MRI dataset with multiparametric maps (MPMs) (10 participants; McColgan et al., 2021), and a multimodal dataset with resting-state MEG and 7T MPMs (31 participants). Color-coded are electrophysiological variables (ephys), cytoarchitectonic variables (cyto), and myelin and iron estimates from high-resolution MRI (myelo). FA - factor analysis.

| Step | Goal | Modality | Datasets | Inputs | Preregistered hypotheses | Outcome | Comment |
| --- | --- | --- | --- | --- | --- | --- | --- |
| 1 | Comparing the regional distributions of ephys between EEG and MEG | across regions | LEMON (EEG), Cam-CAN (MEG) | theta power, alpha power, alpha frequency, low beta power, high beta power, 1/f slope estimated in the 2–40 Hz range, 1/f slope estimated in the 40–60 Hz range, nonsinusoidality of alpha rhythm amplitude, non-zero-mean of alpha amplitude | Electrophysiological variables correlate between EEG and MEG | only alpha power, low beta, and high beta correlated with $r > 0.60$ | |
| 2 | Testing directional hypotheses about the relation of ephys with cyto | across regions | LEMON (EEG), Cam-CAN (MEG), BigBrain | alpha power, thickness of layer II, thickness of layer IV, thickness of layer V | alpha power is positively correlated with thickness of layer II, alpha power is positively correlated with thickness of layer IV, alpha power is positively correlated with thickness of layer V, | positive correlation of alpha power with the thickness of layer IV | Based on previous literature (see Table S1), we preregistered directional hypotheses concerning alpha power, alpha frequency, theta power, and the 1/f slope (seven hypotheses). However, we tested only those variables that showed a significant effect in step 1 (three hypotheses) |
| 3 | Testing all-to-all relationships between ephys, cyto, and myelo | across regions | LEMON (EEG), Cam-CAN (MEG), BigBrain, 0.5 mm MRI <sup>1</sup> | alpha power, low beta power, high beta power, supragranular thickness (sum of layers I, II, and III), thickness of layer IV, infragranular thickness (sum of layers V and VI), mean of the staining intensity profile, skewness of the staining intensity profile, superficial myelin component transformed with FA from R1 and R2*, mid-cortical myelin FA component from R1 and R2*, deep myelin FA component from R1 and R2* | - | alpha power positively correlated with the thickness of layer IV, the thickness of layer IV positively correlated with mid-cortical component R2*, mid-cortical component R2* positively correlated with alpha power | We preregistered this exploratory analysis |
| 4 | Testing relationships between ephys and myelo | across participants | Multimodal dataset (MEG and 0.65 mm MRI) | alpha power, mid-cortical FA component R2* | alpha power positively correlates with mid-cortical FA component R2* | alpha power correlates negatively with mid-cortical FA component R2* | Based on the results of step 3, we preregistered the single directional hypothesis that alpha power correlates positively with mid-cortical FA component R2*. |
| 5 | Testing all-to-all relationships between ephys and myelo | across participants | Multimodal dataset (MEG and 0.65 mm MRI) | alpha power, superficial myelin FA component from R1 and R2*, mid-cortical myelin FA component from R1, deep myelin FA component from R1 and R2* | - | alpha power correlated positively with superficial myelin FA component R1 | We preregistered this exploratory analysis. |

<sup>1</sup> 7T MRI dataset with a resolution of 0.5 mm was previously collected for another study (McColgan et al., 2021)

The depth surfaces of myelin-sensitive multiparametric map (R1 and R2*) were highly correlated with each other. To reduce the correlation and the number of corrections for multiple comparisons, we transformed sampled surfaces into representative components using a factor analysis (FA; see Fig. 2 for all components). FA components were selected based on correlations with depth surfaces (peak correlation higher than 0.3) and are named hereafter with respect to the approximate depth surface with the highest correlation. For example, if the FA component is called “mid-cortical”, it means that this component demonstrated the highest correlation around 50% cortical depth (Fig. 2 and Fig. S2). We used FA on both MRI datasets (see Table 1). Corresponding factors were significantly correlated between two datasets (superficial R1 regression dataset 1 to dataset 2 across vertices: beta = 0.17, *t*-value = 13.62, significant p < 0.001 based on permutations with spins; mid-cortical R1: beta = 0.62, *t*-value = 63.80, *p* < 0.001; deep R1: beta = 0.17, *t*-value = 12.97, *p* < 0.001; superficial R2*: beta = 0.44, *t*-value = 41.61, *p* < 0.001; mid-cortical R2*: beta = 0.74, *t*-value = 91.49, *p* < 0.001; deep R2*: beta = 0.14, *t*-value = 11.98, *p* < 0.001).

For directional hypotheses (see Table 1, step 2), we used thickness values from BigBrain of separate layers (layer II, layer IV, and layer V). For all-to-all correlations (see Table 1, step 3), we combined the thickness of layer I, layer II, and layer III into supragranular thickness by summing the values to reduce the number of multiple comparisons. Similarly, infragranular thickness was the sum of the thickness of layer V and layer VI. Additionally, we tested the staining profile derivatives: profile mean, and profile skewness (distributions on Fig. 2). Staining profile mean reflects the total amount of cell bodies stained in a vertex (in a staining section), and staining profile skewness shows the distribution of cell bodies, with negative skewness corresponding to a higher amount of cell bodies in infragranular layers and skewness around zero corresponding to a relatively symmetrical distribution with a peak at mid-cortical depth.

### Between regions, alpha power correlates positively with thickness of layer IV as well as with mid-cortical R2*

We preregistered three directional hypotheses based on previous literature (Table 1, step 2; Fig. 3a). We used “spin” null models (Váša et al., 2022; Hansen et al., 2022) to control for the spatial smoothness in the distribution of all parameters. Then, we derived *p*-values as a fraction of *t*-values from regressions using spinned data that exceeded the original *t*-value. Additionally, we used bootstrapping to check whether a subset of data points would still support the observed effect. The analysis was performed on vertex and parcellated data (with Julich-Brain Atlas 3.1; Amunts et al., 2020; Amunts et al., 2023).

**Figure 3.**
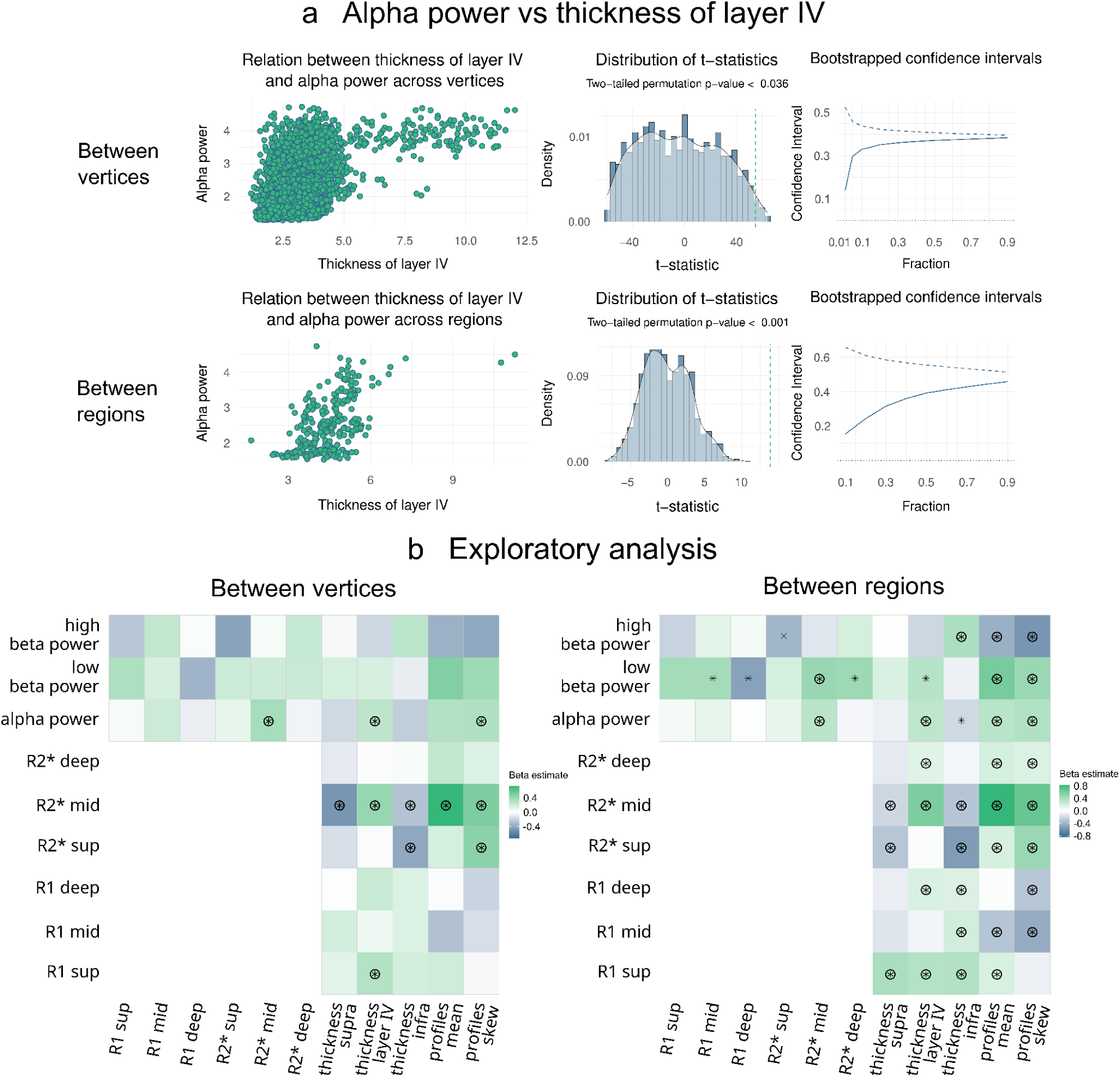
Alpha power correlates with the thickness of layer IV between vertices and regions, as well as with myelin content. **a**. Alpha power and thickness of layer IV are positively correlated. The data were standardized by dividing by their standard deviation. Significance was estimated based on permutation tests with spins. **b**. Associations between three modalities (electrophysiology, cytoarchitecture, and *in vivo* myeloarchitecture) were assessed. Each regression outcome was tested for significance with spin tests and corrected for multiple comparisons with the false discovery rate method (FDR; Benjamini–Hochberg method; Benjamini et al., 1995). Each significant regression coefficient was subjected to a bootstrap procedure. ⊛ = the relation was significant, and the bootstrapped confidence intervals did not exceed zero; ✴ = the relation was significant based on a permutation test, but the confidence intervals crossed zero (implying that the correlation was driven by a subset of vertices/regions). R1 sup - R1 superficial, R1 mid - R1 mid-cortical, R2* sup - R2* superficial, R2* mid - R2* mid-cortical, thickness supra - thickness supragranular, thickness infra - thickness infragranular, profile skew - profile skewness.

Our data only supported the second hypothesis (Fig. 3ab)—alpha power correlated positively with thickness of layer IV (between vertices: beta 0.4, *t*-value = 54.88, 95% CI [0.38, 0.4], two-tailed permutation *p*-value < 0.036; between regions: beta 0.49, *t*-value = 13.89, 95% CI [0.42, 0.56], two-tailed permutation *p*-value < 0.001). Alpha power did not correlate significantly with the thickness of layer II (between vertices: beta –0.16, *t*-value = –21.33, 95% CI [–0.17, –0.14], two-tailed permutation *p*-value = 0.37; between regions: beta –0.06, *t*-value = –1.74, 95% CI [–0.13, 0.01], two-tailed permutation *p*-value = 0.52). Alpha power correlated significantly but negatively with the thickness of layer V (between vertices: beta –0.30, *t*-value = –41.65, 95% CI [–0.32, –0.29], two-tailed permutation *p*-value < 0.03; between regions: beta –0.24, *t*-value = –6.592, 95% CI [–0.32, –0.17], two-tailed permutation *p*-value < 0.002).

In addition to our directional hypotheses, we also performed analyses for all variables (Table 1, step 3). We corrected for multiple comparisons using the false discovery rate method (FDR; Benjamini–Hochberg method; Benjamini et al., 1995). Significance was evaluated with permutation testing using spins. To make sure that the associations were not driven by a few vertices (or regions), we performed bootstrapping (see Methods). To this end, we ran the same correlations but randomly sampled a subset from the whole pool of vertices (or regions) taken for the correlation (the lowest fraction for vertex-based analysis was 1% which corresponds to 90 vertices; the lowest fraction for region-based analysis was 10% which corresponds to 27 regions).

Between the three modalities, we found converging correlations (Fig. 3c): alpha power correlated significantly with mid-cortical R2* (between vertices: beta 0.36, *t*-value = 38.43; between regions: beta 0.32, *t*-value = 6.79), which in turn correlated significantly with cytoarchitectonic variables, including thickness of layer IV (between vertices: beta 0.42, *t*-value = 40.37; between regions: beta 0.59, *t*-value = 13.03). Both alpha power and mid-cortical R2* correlated with profile skewness as well. R2* mid-cortical correlated positively with profiles mean (which is an approximation of the total amount of cells), but the correlation of alpha power and profiles mean did not reach significance (see Fig. S4 for the same analysis of the Multimodal dataset).

### Between participants, alpha power correlates negatively with mid-cortical R2*

Since the results for the between-region analysis demonstrated a correlation of alpha power and mid-cortical R2*, we tested the same hypothesis between participants (Table 1, step 4). For each vertex (or each region), we estimated the relationship between alpha power and mid-cortical R2* (as a measure of myelin and iron). Additional confounds in the regression model were age, brain volume, skull thickness, and local curvature. In the vertex-wise analysis, we observed a significant negative correlation between alpha power and myelin estimates. Thus, participants with larger mid-cortical R2* exhibited lower alpha power at rest, compared to those with lower levels of myelin. This association was particularly pronounced around central and frontal regions (pre-central gyrus, post-central gyrus, superior parietal lobule, superior frontal gyrus, Fig. 4). Two significant clusters were revealed with permutation testing (average beta in clusters = –0.29; average *t*-value = –2.59). A similar correlation was observed for 50%-depth R2* surface (Fig. S5). In the region-based analysis, 23 regions survived the multiple comparison correction with FDR (average beta in significant regions = –0.36; average *t*-value = –3.53; Fig. 4; see Table 2 for the list of regions; the parcellation used was Julich Brain Atlas 3.1; Amunts et al., 2020; Amunts et al., 2023). One region showed a positive correlation (beta = 0.30, *t*-value = 3.15). Additionally, we tested the correlation between other myelin estimates and alpha power (Table 1, step 5). No other component of R2* or R1 demonstrated significant correlations in the vertex-based analysis. However, in the region-based analysis, we found two regions in the superficial R1 component (see Results/Selection of variables for definition) that had positive correlation (average beta in significant regions = 0.44; average *t*-value = 4.96; Fig. S6; see Table 2 for the list of regions).

**Figure 4.**
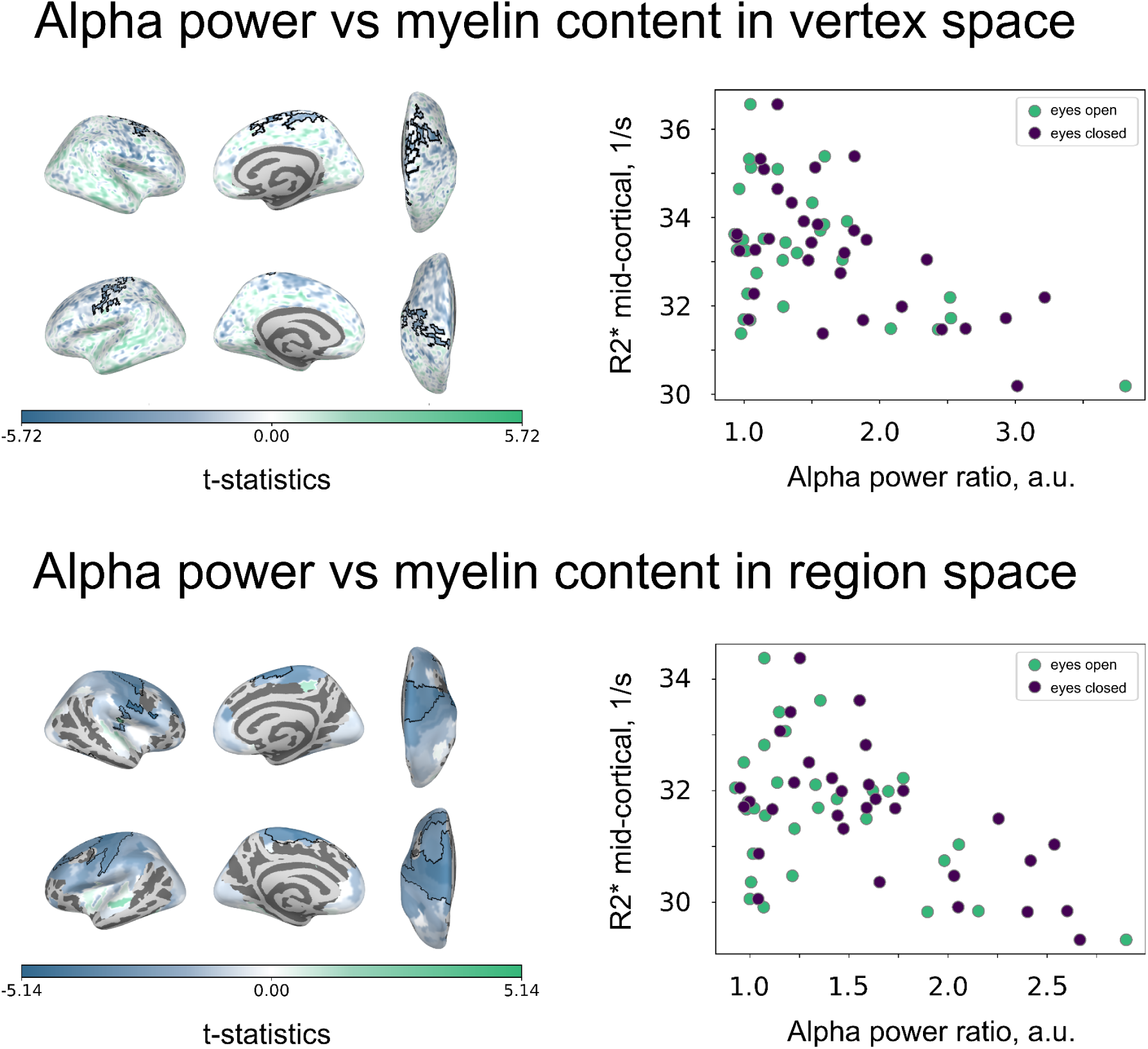
Alpha power correlates negatively with mid-cortical R2* between participants. A significant cluster in vertex space was determined based on the permutation test. Cluster-forming threshold *p*-value < 0.05, cluster significance threshold *p*-value < 0.05 (two-tailed). Significant regions in region space were FDR corrected (corrected *p*-value < 0.05). Each point indicates one participant (averaged within significant clusters or across significant areas with negative correlation).

**Table 2.** The list of regions (from region-based analysis) with significant correlations between alpha power and multiparametric maps (MPMs). Only regions that survived correction for multiple comparisons are shown.

| | Superficial $R1$ | Middle $R2^*$ |
| --- | --- | --- |
| positive correlation with alpha power | post-central gyrus area 3b on the right,<br>superior temporal gyrus area Te 2.2 on the right | parietal operculum subdivision 3 on the right |
| negative correlation with alpha power |  | post-central gyrus area 3a and 3b on the left,<br>pre-central gyrus area 4a on the left,<br>area 4p on the left,<br>area 6d1 bilateral,<br>area 6d2 bilateral,<br>area 6d3 on the right,<br>area 6v1 bilateral,<br>area 6v2 on the left,<br>area 6v3 on the right,<br>superior frontal gyrus area 8d1 on the left,<br>area 8v2 on the left,<br>superior frontal gyrus subdivision 2 and 4 on the left,<br>middle frontal gyrus subdivision 1 on the right,<br>inferior frontal junction on the right,<br>fusiform gyrus subdivision 4 on the right,<br>insula subdivision 2 on the right,<br>parietal operculum subdivision 4 on the right |

### Computational modeling of different effects of excitatory pyramidal neurons and inhibitory interneurons on alpha power

The between-region analysis revealed a positive correlation between alpha power and R2* (index of myelination and iron content) at the mid-cortical depth, a positive correlation between alpha power and the thickness of layer IV, and a positive correlation between mid-cortical R2* and the thickness of layer IV. Yet, from the analysis between subjects, we obtained a negative correlation between alpha power and mid-cortical R2*. To further investigate these discrepancies, we ran an exploratory analysis (not preregistered).

In addition to correlating with the thickness of layer IV, alpha power also correlated (significantly between regions and non-significantly between vertices) with the staining profile mean. The mean of the staining profile provides an approximation for the total amount of cell bodies in one vertex (or in one region). We ran an exploratory mediation analysis to estimate how much of the correlation between the thickness of layer IV and alpha power can be explained via correlation with the profile mean. While the total effect was significant, the direct effect was much smaller. For vertices, 88% of the correlation between the thickness of layer IV and alpha power was explained by the effect of mediation via profile mean (Average Causal Mediated Effect (ACME) beta = 0.27, *p*-value < 0.001; Average Direct Effect (ADE) beta = 0.04, *p*-value = 0.002; total effect beta = 0.30, *p*-value < 0.001; proportion mediated 0.88, *p*-value < 0.001). For regions, the direct effect even changed the direction of correlation (ACME beta = 0.43, *p*-value < 0.001; ADE beta = –0.03, *p*-value = 0.68; total effect beta = 0.40, *p*-value < 0.001; proportion mediated 1.07, *p*-value < 0.001). Thus, the total amount of cell bodies in the staining section (of which the majority are pyramidal neurons; Shepherd & Grillner, 2017) correlates positively with alpha power and substantially explains the correlation between alpha power and the thickness of layer IV between regions. Similarly, correlation between mid-cortical R2* with thickness of layer IV was partially explained by profile mean (for vertices, ACME beta = 0.20, *p*-value < 0.001; ADE beta = 0.15, *p*-value < 0.001; total effect beta = 0.35, *p*-value < 0.001; proportion mediated 0.57, *p*-value < 0.001; for regions, ACME beta = 0.40, *p*-value < 0.001; ADE beta = 0.10, *p*-value = 0.42; total effect beta = 0.49, *p*-value < 0.001; proportion mediated 0.79, *p*-value < 0.001).

Both staining profile mean and myelin estimates incorporate excitatory and inhibitory cells. Yet, we obtained opposing correlations in between-region and between-participant analyses. Therefore, we evaluated the possible contribution of excitatory and inhibitory neurons to alpha power using exploratory simulations with the Wilson–Cowan neural mass model (Wilson & Cowan, 1972) within the Virtual Brain simulation framework (Sanz Leon et al., 2013; Schirner et al., 2022). The motivation to use the Wilson–Cowan neural mass model was that, in this model, parameters for excitatory and inhibitory populations can be tuned separately. For each region of the Desikan–Killiany atlas (Desikan et al., 2006), time series with alpha oscillations were simulated (Fig. 5ab). To create variance in the power of alpha oscillations (Fig. 5c) over regions, we systematically varied two parameters of the excitatory population: the excitatory-to-excitatory coupling coefficient *C_ee_* and the value of the maximum slope of the sigmoid function *a_e_* (Fig. 5d). To create variance across simulated subjects, we varied the external stimulus to the inhibitory population *Q*. We observed that with a decrease in excitatory population coupling and reactivity to the stimulus, alpha rhythm decreased in amplitude (Fig. 5d). This is in agreement with evidence that AMPA and NMDA blockers decrease alpha power (Muthukumaraswamy et al., 2015; Lozano-Soldevilla, 2018). At the same time, with an increase in the input to the inhibitory population, alpha power decreased (Fig. 5d). This is in agreement with evidence that pharmacologically increased inhibition leads to a decrease in the alpha amplitude (Schreckenberger et al., 2004; Ahveninen et al., 2007; Lozano-Soldevilla et al., 2014; Opie et al., 2025 *bioRxiv*). Thus, we infer that the relations that we have observed in our data could be explained with contributions of different types of cells. First, the positive correlation between the thickness of layer IV and total amount of cells with alpha power across regions could be explained by the positive correlation between excitatory activity and alpha power. Second, the negative correlation between myelin estimates and alpha power across participants could be explained by the negative correlation between inhibitory activity and alpha power.

**Figure 5.**
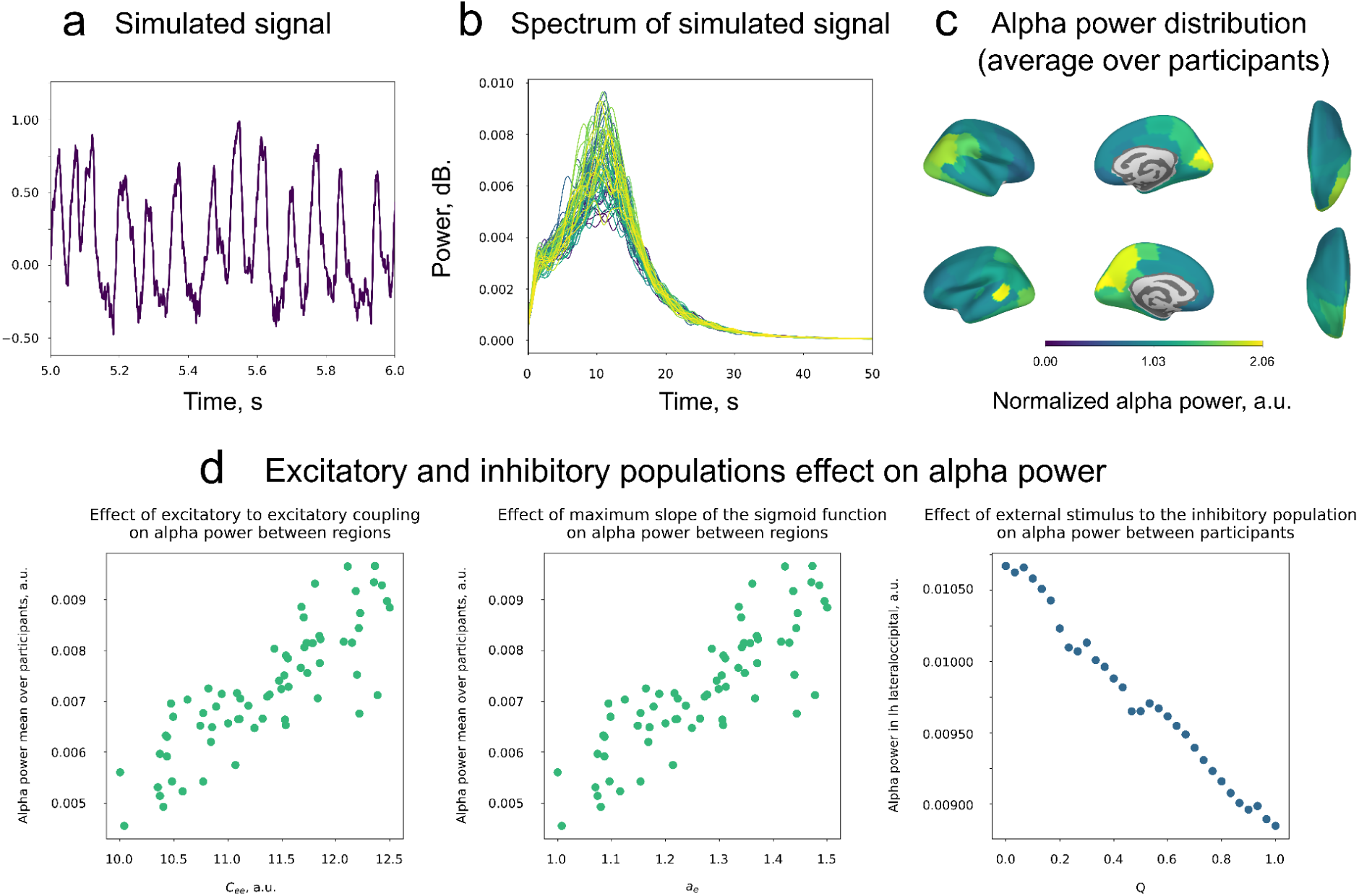
Alpha power may be differentially related to excitatory and inhibitory neurons. **a**. Alpha oscillations were simulated with the Wilson–Cowan model using The Virtual Brain framework. The time series was simulated to resemble periodic oscillations with a 10 Hz frequency. **b.** The spectra of simulated time series revealed the peak at around 10 Hz. **c**. The alpha power was simulated with a typical distribution with larger power in occipital regions. This was achieved by varying the parameters of the excitatory population: the excitatory-to-excitatory coupling coefficient *C_ee_* and the value of the maximum slope of the sigmoid function *a_e_* (which defines mapping from external input to firing rate of a population). **d**. Regional differences in alpha power were simulated by varying parameters of the excitatory population, whereas inter-individual differences were modeled by adjusting the external input to the inhibitory population (*Q*). The increase in activity of the excitatory population led to an increase in alpha power, while an increase in activity of the inhibitory population led to a decrease in alpha power.

## Discussion

We assessed how functional electrophysiological variables relate to microstructural features of the cortex by studying their variation both across cortical regions and across individuals. To this end, we combined open-access high-resolution histological data, large-cohort EEG/MEG and MRI datasets, and a newly acquired dataset combining structural 7T MRI data and MEG data collected from the same group of participants. Our analyses revealed a cross-regional pattern: alpha power increased with the thickness of layer IV, which in turn increased with mid-cortical myelin and iron as estimated by R2*. Moreover, alpha power increased with mid-cortical R2*. In other words, regions with a thicker layer IV tended to exhibit higher myelin and, consequently, higher alpha power. In contrast, across participants, we observed the opposite relationship: a higher mid-cortical R2* was associated with lower alpha power. We attribute the difference in correlation sign to the distinct regional dominance of excitatory and inhibitory processes across the brain, which differentially shape the observed relationships. We performed simulations with a mass model to support this explanation.

### Selected variables

Since our goal was to link laminar microstructure to macroscopic rhythms rather than to exhaustively screen diverse spectral metrics, the analyses were restricted to electrophysiological variables with robust and reproducible spatial distributions across large EEG and MEG cohorts. Only alpha, low beta and high beta power met the preregistered criterion of strong EEG–MEG topographic agreement (correlation of >0.6). Subsequent structure–function analyses focused on these bands. Alpha rhythm emerged as the only rhythm that consistently related to cytoarchitecture and depth-resolved myeloarchitecture.

We applied dimensionality reduction with factor analysis to the structural MRI data, motivated by the observation that depth surfaces were strongly correlated with each other. This occurred partially due to resolution (our MRI data were of 0.5 mm and 0.65 mm), which, with respect to cortical thickness, is still quite coarse, as cortical thickness in some regions reaches only 1.5 mm. To mitigate this correlation, we used factor analysis and obtained three distinct factors that were easily interpretable with respect to initial depth surfaces and were remarkably similar between the two datasets. While intracortical myelin includes two types of fibers: tangential and radial fibers (Geyer & Turner, 2013), factor analysis did not accomplish the separation of the two types. The mid-cortical factor still seems to reflect both fiber types, as mid-cortical R2* correlates with staining profile mean (total cell amount) and with supragranular, granular, and infragranular thickness. In general, there were no one-to-one correlations of factors to thickness values or depth-specific myelin estimates (e.g., superficial R1 correlated with thickness of layer IV; while the mid-cortical factor peaked at middle layers, the deep factor also showed a significant loading at those depths). Therefore, one should be cautious when interpreting the results in a layer-specific manner. Nonetheless, we suggest that factor analysis for high-resolution MRI data may be an effective approach for decorrelating the data and for dimensionality reduction.

We observed significant correlations mostly with R2*, while R1 did not demonstrate such results. While R2* is sensitive to myelin and iron content, R1 is mostly sensitive to myelin (Edwards et al., 2018). In a previous study (McColgan et al., 2021), significant correlations with cell numbers and 7T MRI myelin estimates were also observed only for R2* but not for R1 (we used the same dataset for across-region analysis). This may be because R2* has a higher signal-to-noise ratio at 7T (Weiskopf et al., 2014; Tabelow et al., 2019), and because precise R1 mapping is, unlike R2* mapping, also dependent on the quality of transmit B1 mapping (Edwards et al., 2023). Additionally, R2* contains a partial contribution from blood hemodynamics (McColgan et al., 2021). However, it is unlikely that blood flow could explain our results. Firstly, the density of vessels increases towards the pial surface (Havlicek & Uludağ, 2020; Haenelt et al., 2025), and we did not find a correlation between alpha power and superficial R2*. Even if those vessels penetrate the cortical mantle, we would expect to find similar correlations for superficial, mid-cortical, and deep R2*, but we did not. Also, while blood flow relates to metabolic activity (Raichle et al., 2006), the relation of metabolic activity to cell density is not established (Ventura-Antunes et al., 2022). Secondly, alpha power was previously associated with blood perfusion in the brain (O’Gorman et al., 2013): participants with higher perfusion (i.e., increased blood flow) also had higher alpha power. If our results were driven simply by blood volume, then larger blood volume (and larger signal in MRI) would lead to larger alpha power, which was not the case.

### Intra-individual structure-function coupling

Among electrophysiological variables (as measured with large LEMON and Cam-CAN datasets), only alpha power showed robust correlations both with cytoarchitectonic variables (namely, positive correlation with thickness of layer IV and staining profile skewness) and with high-resolution MRI (namely, positive correlation with mid-cortical R2*). Namely, regions with a thicker layer IV also had more myelin and iron and greater alpha power. Layer IV, or the granular layer, is a main entry layer for signals from the sensory thalamus (Shepherd et al., 2017), and alpha power positively correlates with thalamic activity (Goldman et al., 2002; Moosmann et al., 2003; Schreckenberger et al., 2004). The granular layer contains both inhibitory interneurons and excitatory pyramidal neurons (Scala et al., 2019; Jorstad et al., 2023). Therefore, it could be that the presence of more pyramidal neurons within a thicker layer IV may facilitate stronger alpha rhythm generation. However, we also observed that the thickness of layer IV was highly correlated with the mean of the staining profile, which serves as an approximation of total cell amount. That indicates that the mean of the staining profile explained almost all of the correlation between the thickness of layer IV and alpha power between vertices (and regions). This suggests that alpha power in a region may be determined by the total number of neurons in that region. This, in turn, would align well with earlier predictions that stronger oscillation amplitudes are likely to reflect a larger number of participating neurons (Pfurtscheller et al., 1999). In the same way, myelin and iron content as captured by mid-cortical R2* may also be to a large extent influenced by the total number of neurons. We thus propose that intra-individual variability can be explained by the total number of neurons, which lead to more axons, more myelin, and more alpha.

### Inter-individual structure-function coupling

Across participants, the same mid-cortical R2* measure that related to alpha positively across regions, related negatively to alpha power, most prominently in somatomotor and frontal cortex, with a broadly negative trend across the cortical mantle. The detection of this significant correlation was most likely facilitated by better data quality of MPMs (see Fig. S3 for error maps; Mohammadi et al., 2022). The correlation pattern did not follow alpha power distribution, suggesting that it was not driven by signal-to-noise ratio in the alpha band. The opposite direction of correlation implies that inter-individual and intra-individual differences must be influenced by different factors.

The difference between the signs of these correlations suggests that some cytoarchitectonic or functional characteristics contribute differently to the variability between individuals (inter-individual) and between regions (intra-individual). Previously, it has been shown that the difference in cytoarchitecture is larger between individuals than between two regions of the language network, Brodmann’s areas 44 and 45 (Amunts et al., 1999). Moreover, the variance of neurotransmitter receptors, including GABA and glutamate, is larger between people than between regions (Hansen et al., 2026). Further, alpha power and GABA and glutamate concentrations correlate oppositely between and within individuals (Cochrane et al., 2025 bioRxiv). Possibly, variable trends can be explained by microcircuits involving excitatory and inhibitory neurons.

Intracortical myelin primarily includes two types of fibers: tangential and radial fibers (Geyer & Turner, 2013). While radial fibers most likely originate from ascending and descending axons, tangential fibers are thought to largely represent myelinated axon collaterals of pyramidal neurons (Hellwig, 1993) and short-range axons of interneurons (Stedehouder et al., 2017; Micheva et al., 2021). Intuitively one might think that if there are more incoming myelinated radial fibers, then the input to the cortex would be more synchronous, which would lead to larger alpha power. However, that is not what we observed in the analysis across participants. If there were more departing myelinated fibers, the effect would be expected in the deep layers. However, we observed no significant correlation of alpha power with either deep R1 or deep R2*. Therefore, these differences across participants can rather be explained by tangential fibers.

Tangential fibers that are collateral axons of pyramidal neurons may make synapses with inhibitory neurons downstream (Berger et al., 2010). This would result in increasing inhibition in the vicinity of pyramidal neurons. Instead of having a larger cortical patch producing oscillations in synchrony, lateral inhibition will create several smaller patches that are not synchronized with each other (Kumral et al., 2022). As a result, on the scalp, we would observe a smaller alpha rhythm amplitude. Furthermore, tangential fibers that are axons of inhibitory neurons may indicate the presence of more inhibitory neurons and hence more inhibition. Pyramidal neurons, while being inhibited, do not lose the rhythm but rather produce the rhythm with a smaller amplitude (Antkowiak, 1999; Lozano-Soldevilla et al., 2014). We thus propose that inter-individual variability can be explained by the difference in inhibitory modulation.

Previous research showed that variability of cytoarchitecture and neurotransmission between regions and between participants is indeed different (Amunts et al., 1999; Cochrane et al., 2025 bioRxiv; Hansen et al., 2026), but the causes remain unknown. Our explanation rests on the assumption that it is variability of excitatory and inhibitory neurons and their fibers that is different between regions and between participants. Even though inhibitory neurons comprise only 20–30% of all neurons (Shepherd & Grillner, 2017), if variability is larger in comparison to variability of excitatory neurons, even a small amount can produce large-scale effects (which can be recorded with non-invasive recordings). Future studies are needed to elucidate the microstructural (e.g., cytoarchitectonic) influences of intra-individual and inter-individual variability.

## Conclusion

In the current study, we identified an association between electrophysiological variables and cyto- and myelo-architecture. Resting-state alpha power covaried with cortical microstructure in two systematically different ways. Across regions, alpha was higher in areas with a thicker layer IV and greater mid-cortical myelin and iron estimates (R2*). Whereas across individuals, in frontal and central cortex, alpha was lower in participants with higher mid-cortical R2* component. We propose one possible explanation via the effects of excitatory pyramidal neurons versus the effects of inhibitory interneurons. In particular, positive correlation across regions may be related to the total number of cells in the region, while negative correlation across participants—to the difference in inhibition that varies between participants. Further research should focus on determining the structural and functional factors that contribute to intra-individual and inter-individual differences.

## Methods

The study was preregistered (Studenova, 2023; see also Table 1).

### Datasets

#### EEG LEMON dataset

The LEMON dataset (Babayan et al., 2019) contains EEG data from 216 participants (209 after preprocessing), aged 20–77 years old. The data was collected at the Day Clinic for Cognitive Neurology of the University Clinic Leipzig and the Max Planck Institute for Human Cognitive and Brain Sciences (MPI CBS) in Leipzig, Germany. Participants were preselected based on medical and psychological screening and considered to be healthy (see Table 1 in Babayan et al., 2019). The EEG was recorded with a BrainAmp MR plus amplifier with 61 scalp electrodes and a single VEOG channel. EEG electrodes were active ActiCAP electrodes placed on the scalp according to the standard 10–20 localization system. The reference electrode was FCz. The ground electrode was placed at the sternum. The recording was performed with a bandpass filter between 0.015 Hz and 1 kHz and a sampling rate of 2500 Hz. The resting session contained 16 blocks of one-minute duration, with eyes-open and eyes-closed blocks interleaved (i.e., 8 minutes total for each condition). Participants were asked to stay awake and, during the eyes-open condition, fixate on the cross on the screen. We included all available data.

#### MEG Cam-CAN dataset

The Cambridge Centre for Ageing and Neuroscience dataset (Cam-CAN; Shafto et al., 2014; Taylor et al., 2017) was acquired and shared by the Cambridge Centre for Ageing and Neuroscience (Cam-CAN). Funding for Cam-CAN was granted by the UK Biotechnology and Biological Sciences Research Council (grant number BB/H008217/1), with additional support from the UK Medical Research Council and the University of Cambridge, UK. The dataset contains the MEG, MRI, and fMRI data of 572 participants, aged 18–87 years old. We excluded participants if they were diagnosed with one of the following conditions: stroke, meningitis or encephalitis, insomnia requiring treatment, Parkinson’s disease, motor neuron disease, multiple sclerosis, epilepsy, head injury, brain operation, brain tumor, bipolar disorder, schizophrenia, Tourette syndrome, dementia/Alzheimer’s disease, or drug abuse. The MEG was recorded with the MEG system VectorView Elekta Neuromag, 102 magnetometers, and 204 orthogonal planar gradiometers in a magnetically shielded room. For our purposes, we used MEG resting-state recording with eyes closed. The duration of the data was at least 8 minutes and 40 seconds for each participant. The data was recorded with a sampling rate of 1000 Hz and a high-pass filter of 0.03 Hz. We had no exclusion criteria based on data quality. The data were sourced from the Cam-CAN repository (http://www.mrc-cbu.cam.ac.uk/datasets/camcan; Shafto et al., 2015; Taylor et al., 2017).

#### BigBrain and Julich Brain Atlas

We used BigBrain (Amunts et al., 2013) to obtain cytoarchitectural variables. BigBrain is a free and publicly available ultrahigh-resolution model of the postmortem paraffin-embedded brain of a 65-year-old male. The brain was cut into 7404 histological sections, each of 20 μm thickness, which were then stained for cell bodies and digitized. The sections were reconstructed into a 3D model (Amunts et al., 2013).

For the region-based analysis, we used the Julich-Brain Atlas v3.1 (JBA; Amunts et al., 2020; Amunts et al., 2023). This atlas includes data from ten postmortem brains, thus carrying information about intersubject variability. The parcels in the atlas are probability maps of cytoarchitectonically defined regions over the cortical surface and subcortical nuclei. Thus, each voxel has an associated probability of belonging to a particular region based on the analysis of the aforementioned brains. For parcellation, we used maximum probability maps, where each voxel is assigned to the particular region when it is the most probable assignment.

#### 7T MRI dataset (I)

For myelin estimates, we used the dataset from McColgan et al. (2021). Briefly, the dataset comprised 7T MRI scans with an isotropic resolution of 0.5 mm acquired from 10 healthy participants (6 females, 22–33 years old). The protocols consisted of two multi-echo fast low angle shot (FLASH) scans with T1- and PD-weighting (T1w, PDw) along with maps of the radio frequency (RF) transmit field B1+ and static magnetic field B0. These images were then used to estimate R1 and R2* parameter maps using the hMRI toolbox (Tabelow et al., 2019). We used the preprocessed data from the original study.

#### Multimodal 7T MRI (II) and MEG dataset

For microstructural estimates, we recorded 7T structural MRIfrom 31 participants (15 females, 21–35 years old) using well-established MRI protocols (Marques et al., 2010; Trampel et al., 2019; Kirilina et al., 2020; McColgan et al., 2021). All participants signed written informed consent. The study was approved by the ethics committee of the University of Leipzig (318/24-ek). We used a 7T whole-body MRI system (Siemens Magnetom 7.0T W60 Numaris/X VA60A-0CT2, Siemens Healthineers, Erlangen, Germany) equipped with an 8-channel transmit/32-channel radio-frequency (RF) receive head coil (Nova Medical, Wilmington, MA). The protocol consisted of three multi-echo fast low angle shot (FLASH) scans with T1-, PD- and magnetization transfer- (MT-) weighting and 3D-EPI spin echo/stimulated echo data to calculate maps of the radio frequency transmit field B1+ for flip angle correction. Acquisition parameters were: readout bandwidth of 445 Hz/pixel; TR=22.4 ms; TEs between 3 ms and 15.6 ms (△TE = 2.52 ms, 6 echos) for T1w and PDw and between 3 ms and 10.56 ms (△TE = 2.52 ms, 4 echos) for MT-weighted; excitation flip angles: 7° (PDw and MTw), 22° (T1w); MT pulse Gaussian at 3 kHz offset, with flip angle 130° and duration 4 ms. Additional parameters for each weighting: isotropic voxel size 0.6×0.6×0.6 mm³, CAIPIRINHA acceleration factor 2×2 (Breuer et al., 2006), acquisition time 8:25 min per contrast. These images were used to estimate the R1 and R2* parameter maps with the use of the hMRI toolbox v0.6.1-dev git hash 863fa08 (Tabelow et al., 2019). For segmentation purposes, we obtained an MP2RAGE acquisition (Marques et al., 2010) with the following parameters: voxel size 0.8×0.8×0.8 mm³, slice thickness 0.75 mm, TR 4300.0 ms, and TE 2.27 ms.

For electrophysiological estimates, we recorded MEG with the VectorView Elekta Neuromag system with 102 magnetometers, and 204 orthogonal planar gradiometers in a magnetically shielded room. Participants were asked to relax and not think about anything in particular. In total, the data contained 16 blocks, each 60 seconds long, with open- and closed-eyes blocks interleaved (eight open-eyes blocks and eight closed-eyes blocks), starting from an open-eyes block. The transition between blocks was signaled by a sound signal. During the open-eyes block, the participants were asked to fixate their eyes on the black cross on the screen. The data was recorded with a sampling rate of 2500 Hz and a low-pass filter of 750 Hz was applied. In addition, EOG (vertical and horizontal) and ECG signals were recorded for later artifact cleaning.

The average break between two sessions was seven days (maximum 24 days, minimum 1 day, average 7 days). Seven participants had the MEG session first, while 24 participants had the MRI session first. The MEG session never occured the day after the MRI session.

### Preprocessing

#### EEG data

The preprocessing of EEG data was performed with the MNE-Python package (Gramfort et al., 2013). For each participant, we followed a fairly standard pipeline. After loading the data, we removed noisy channels based on visual inspection, then re-referenced the data to a common average reference (Osselton, 1965). Then, we filtered the recording in a wide bandpass range, from 0.1 Hz to 100 Hz, with the addition of a notch filter around 50 and 100 Hz. Afterwards, we split the data into two conditions, eyes-open and eyes-closed. When splitting the data, we removed one second around the splitting triggers to avoid artifacts caused by participants adjusting to a new condition. For both conditions, we removed segments of data with high-amplitude artifacts (detected with autoreject; Jas et al., 2016; Jas et al., 2017). For the eyes-open condition, we ran ICA and discarded two components related to vertical and horizontal eye movements (based on the similarity of the time course with the signal from the VEOG electrode). Lastly, the data was downsampled to a sampling rate of 500 Hz.

The sensor data from the LEMON dataset for each condition were reconstructed into the source space surface. We applied source localization based on the fsaverage subject (obtained with mne-python) from FreeSurfer (Fischl, 2012) and a 3-layer Boundary Element Method (BEM) model to compute the forward model. As a source reconstruction method, we used eLORETA (Pascual-Marqui et al., 2011) with the following parameters: free-orientation in the inverse operator; constrained to the normal of the cortical surface orientation of dipoles; regularization parameter lambda = 0.05; and the covariance of white noise with equal duration to the data as noise covariance (Idaji et al., 2022). The reconstructed surface had 8196 candidate dipole locations in each hemisphere.

#### MEG data

##### Cam-CAN

We used pre-processed data that is available from the Cam-CAN. The detailed preprocessing pipeline is described elsewhere (Taylor et al., 2017). Briefly, the following steps were performed: 1) temporal signal space separation (tSSS, Taulu et al., 2005; MaxFilter 2.2, Elekta Neuromag Oy, Helsinki, Finland) to clean data from external sources and from HPI coil noise, to correct for head-motion, and to transform data to a common head position; 2) 50 Hz notch filter; 3) detection and interpolation of noisy channels; 4) exclusion of ICA components correlated with eye movements and pulse-related artifacts (total of three components).

The sensor data from the Cam-CAN dataset were reconstructed into the fsaverage source space, following the same steps as for the LEMON data, except that the BEM model only had one layer, and noise covariance was derived from empty-room recordings.

##### Multimodal MEG dataset

The preprocessing of MEG data followed the MNE-Python-based FLUX Pipeline (Ferrante et al., 2022) with additional quality assessment steps. 1) bad MEG channels were identified with manual inspection followed by an automated procedure (using mne.preprocessing.find_bad_channels_maxwell). 2) tSSS/SSS and Maxwell filtering was performed. 3) muscle artifacts were automatically annotated. 4) ICA was applied to remove eye blinks, eye movements, and heartbeat-related artifacts: ICA components for rejection were identified with manual inspection and removed from the data. 5) data were downsampled to 1000 Hz and then split into two conditions, eyes-open and eyes-closed, which were later analyzed separately. When splitting the data we removed one second around splitting triggers to avoid artifacts caused by participants adjusting to a new condition.

We performed the same source reconstruction as for the Cam-CAN dataset, except that we used individual participant MRIs for reconstruction with subsequent warping to fsaverage.

#### MRI data

##### 7T MRI dataset (I)

We used pre-processed data. A detailed description of the preprocessing steps can be found in McColgan et al. (2021). In brief, R1 and R2* multiparametric maps were estimated with the hMRI toolbox (Tabelow et al., 2019). Each map was then interpolated to the cortical surface at 10 equivolumetric cortical depths. Creating cortical surfaces was achieved via the following steps: 1) grey matter (GM) and white matter (WM) tissue probability maps were derived with the CAT12 toolbox (http://www.neuro.uni-jena.de/cat); 2) GM and WM were conservatively pruned by setting values below 1 to 0; 3) additional manual correction was applied to improve small segmentation errors; 4) GM and WM probability maps were then used to create 10 equivolume depths using Nighres (Huntenburg et al., 2018).

The obtained depth surfaces were highly correlated with each other, partially due to resolution (0.5 mm; while the thickness of cortex varies from 1 mm to 3.5 mm) and partially due to the ascending and descending axons spanning several cortical layers. Therefore, we applied spatial decomposition with Factor Analysis (sklearn.decomposition.FactorAnalysis; the number of components was equal to 10, rotation was varimax) to reduce dimensionality and obtain depth-specific components for each of the MPMs. Components resulting from the factor analysis were selected based on their correlation with depth surfaces (peak correlation of more than 0.3 and correlation profile favors certain depths, see Fig. S2). Additionally, previous studies using the same dataset demonstrated that R1 maps are affected by inaccuracies of the B1+ field mapping when B1+ becomes low in the temporal and inferior frontal lobes (Edwards et al., 2023). To reduce this confound, for R1 only, we created a mask based on the B1+ map at the mid-cortical surface (threshold B1+ at 60% of nominal). Lastly, the data for each component were smoothed with a 3 mm Gaussian smoothing kernel and downsampled to yield 8196 vertices (to make further computation feasible).

##### 7T MRI dataset (II)

To transform R1 and R2* multiparametric maps to depth surfaces, we first ran FreeSurfer 6.0 recon-all (Fischl et al., 2004) for cortical surface reconstruction. As input to recon-all, we used the MP2RAGE image acquired in the same session. The following steps were performed. 1) background noise was removed using regularization as in O’Brien et al., (2014) (function robust_combination in fMRI tools). 2) bias field correction was applied to the UNI image. 3) the resulting bias-field corrected UNI image was fed into autorecon1 with the *hires* flag without skull stripping. 4) skull stripping was performed in SPM from the second inversion image of the MP2RAGE (the tissue class exclusion thresholds were cerebro-spinal fluid 20%, bone 10%, soft tissue 10%, air 10%). 5) autorecon 2 and 3 were executed. 6) pial and white surfaces were slightly smoothed. 7) resulting pial and white surfaces were coregistered to the MPM image using fMRI tools 1.0.0 (Haenelt, 2022; https://github.com/haenelt/FmriTools), in particular, for registering MP2RAGE surfaces and MPMs, we used the T1w image of the MP2RAGE and R1.

After running the pipeline, we generated 10 equivolume surfaces between the outer (pial) and the inner (white) surfaces. Lastly, R1 and R2* volumes were sampled at each cortical depth. We extracted approximations of myelin at different cortical depths with factor analysis (in the same way as described in an earlier section). Each component was smoothed with a 3 mm Gaussian smoothing kernel.

##### Cytoarchitecture

To obtain thicknesses of each layer, we used the BigBrainWarp toolbox (Lewis et al., 2020; Paquola et al., 2021). The thickness values for each layer were precomputed in BigBrainWarp in the fsaverage reference space. To obtain profile derivatives, we used the staining intensity profiles from Wagstyl et al. (2018). Each profile spans a direction from the pial surface to the white matter border and is defined per vertex in the BigBrain reference space. For each profile (each vertex), we computed the mean value and skewness and then transformed the values into fsaverage space using BigBrainWarp. For smoothing the resulting spatial distributions, we applied the HCP workbench command -metric-smoothing (Marcus et al., 2011). The data for each feature were downsampled to yield 8196 vertices.

### Statistical analysis

#### Variables

For each vertex on the surface of the reconstructed EEG/MEG source space of each participant, we computed the following electrophysiological variables: alpha rhythm power and frequency, theta rhythm power, beta rhythm power, 1/f slope, and alpha rhythm shape. Alpha rhythm power was computed as the power ratio in individual peak frequencies ± 2 Hz to flanking frequencies ± 2–3 Hz. Theta rhythm power was quantified as a ratio of 4–7 Hz to 1–50 Hz. Low beta and high beta rhythm power were quantified as ratios of 15–20 Hz to 1–50 Hz and 20–30 Hz to 1–50 Hz, respectively. The alpha rhythm frequency was computed as a peak frequency in the 7–13 Hz range in the spectra at each dipole location using the find_peaks function (scipy.signal; Virtanen et al., 2020). If multiple peaks were found, the peak with maximal power was selected. The aperiodic component slope (or 1/f slope) was estimated with the FOOOF (fitting oscillations & one over f; Donoghue et al., 2020) algorithm in two ranges: 2–40 Hz (as in Cesnaite et al., 2023) and 40–60 Hz (to avoid oscillatory peaks). For all variables, we computed the spectrum of each vertex with Welch’s method (Welch, 1967; scipy.signal.welch) on 2-second Hanning windows with 50% overlap. Alpha rhythm non-sinusoidality was quantified with the absolute value of deltaCT (Schaworonkow et al., 2019). The non-zero mean of the alpha rhythm was estimated via the absolute value of the baseline-shift index (BSI; Nikulin et al., 2010). The data of all participants were averaged for each vertex and each dataset, namely, the averaging was performed separately for LEMON eyes-open, LEMON eyes-closed, and Cam-CAN. We compared the distribution of group-averaged variables between EEG and MEG across cortical regions (eyes closed). Only variables with distributions that showed a correlation higher than 0.6 (Pearson) between EEG and MEG metrics were further analyzed (this threshold was preregistered).

For each vertex of histological data, we obtained the thickness of each cortical layer, mean staining intensity across layers (which roughly correlates with the total amount of cells in a vertex), and skewness of staining intensity across layers (which correlates with the asymmetry in the amount of cells and cell size in the supragranular layer in comparison with infragranular layers). Thicknesses derived from BigBrain were combined into supragranular (layers I, II, and III) and infragranular (layers V and VI) components by summing respective thickness values.

For each vertex of MRI data, we obtained approximations of myelin and iron as measured with R1 and R2* at three cortical depths (superficial, mid-cortical, and deep) using factor analysis. Factor analysis was performed separately on R1 and R2* maps. Correlation between factors and original depth surfaces is shown on Fig. S2.

#### Between regions

##### Vertex-based analysis

To assess the associations, we used linear mixed-effects modeling. Before model fitting, all variables were standardized by dividing by their standard deviation.

For the directed hypotheses, we fitted a linear mixed-effects model across vertices with one predictor and one dependent variable, with the dataset included as a random factor. For instance, *thickness of layer IV ∼ alpha power + (1|dataset)*, where the dataset is LEMON eyes open, LEMON eyes closed, or Cam-CAN.

For exploratory analysis, we fit a linear mixed-effects model to test the correlation between electrophysiological, cytoarchitectural, and myeloarchitectural variables. For instance, (over all vertices) *thickness of LIV ∼ alpha power + low beta power + high beta power + (1|dataset)*.

The significance of each model was assessed with “spin” null models (Váša et al., 2022; Hansen et al., 2022). That is, for each variable used, we generated 1000 permutations of spinned data, in which the spatial autocorrelation (i.e., smoothness) was preserved. Spins were generated with the netneurotools package in Python (netneurotools.stats.gen_spinsamples; Liu et al., 2025). The significance based on permutation testing was estimated as the proportion of permuted statistics that exceeded the real statistics, correcting for multiple comparisons with the false discovery rate (FDR; Benjamini–Hochberg method; Benjamini et al., 1995).

To ensure that correlation was not driven by just a few vertices, we ran bootstrapping. For each bootstrapping test with a given fraction of vertices, we randomly selected a number of vertices of that fraction and performed the regression only with these vertices. The fractions for vertex-based analysis were from 100% (8196 vertices) down to 1% (81 vertices). We repeated the bootstrapping procedure for each fraction 10000 times.

##### Region-based analysis

Using the 266 regions of interest in the JulichBrain atlas 3.1, we calculated region-wise statistics by averaging the vertex-wise values inside each region. Then, we repeated all regressions as in the vertex-wise analysis. We controlled for significance with permutation testing using spins. The bootstrapping fractions of region-based analysis were from 100% (266 regions) down to 10% (27 regions).

#### Between participants

##### Vertex-based analysis

At every vertex, we computed the association between the variation of MPMs and electrophysiological variables across participants. The analysis was preregistered only for alpha power. For example, (for each vertex) *R2* mid-cortical ∼ alpha power + total brain volume + skull thickness + local curvature + age + (1|condition),* where the condition is eyes open and eyes closed. Thus, for each vertex, we obtained a *t*-value of how strong the association was. Because nearby data points are not independent, we used a cluster-based permutation test to fuse vertices with suprathreshold *t*-values into clusters and test these clusters for significance. The cluster significance was assessed with permutation testing, where, for each vertex, we shuffled the dependent variable across participants. The cluster-forming threshold was *p*-value<0.05, and the cluster significance threshold was *p*-value<0.05 (two-tailed). The confounds that could affect brain dynamics and myelin were total brain volume, skull thickness, local curvature, and age.

##### Region-based analysis

We collapsed data to the JulichBrain atlas 3.1 by averaging values inside each region and repeated the regression for each region. We used FDR for the correction for multiple comparisons (Benjamini–Hochberg method; Benjamini et al., 1995).

### Computational model

To test our hypothesis regarding the differential effects of excitatory and inhibitory populations on alpha power, we used simulations within The Virtual Brain framework (Sanz Leon et al., 2013; Schirner et al., 2022). First, we created alpha rhythm dynamics based on the Wilson–Cowan neural mass model (Wilson & Cowan, 1972) with parameters tuned in such a way that the dynamics would exhibit stable alpha oscillations (*C_ei_* = 15.0, *C_ie_*= 10.0, *C_ii_* = 1.0, *τ_e_* = 10.0 ms, *τ_i_*= 20.0 ms, *a_i_* = 1.0, *b_e_* = 1.0, *b_i_*= 1.0, *θ_e_* = 4.1, *θ_i_* = 3.5, *r_i_*= 0.0 ms, *r_e_* = 0.0 ms, *P* = 1.4).

The brain was simulated as having 68 regions from the Desikan–Killiany atlas (Desikan et al., 2006). To create regional variability in alpha power, we systematically varied the excitatory population parameters: excitatory-to-excitatory coupling coefficient *C_ee_* and value of the maximum slope of the sigmoid function *a_e_*(*C_ee_* was varied in the interval [10.0, 12.5], *a_e_*was varied in the interval [1.0, 1.5]). The parameter ranges were selected such that the alpha rhythm would still be present and frequency precession would be small. Second, we populated the same modelled dynamics across virtual subjects while changing one additional parameter, the external stimulus to the inhibitory population *Q* (which was varied in the interval [0.0, 1.0]).

For each virtual subject, we simulated 60 seconds of data. Then, for each region, the spectrum was computed with Welch’s method (Welch, 1967), and alpha power was estimated at the peak frequency.

## Acknowledgments

We are grateful for the help with data collection to medical technical assistants at MPI CBS: Annett Wiedemann, Domenica Klank, Anke Kummer, Simone Wipper, and Yvonne Wolff-Rosier; as well as to Kamil KILIC. We also thank Niklas Kuegler for help with MRI data management. We are grateful to Sebastian Bludau and Timo Dickscheid for helpful discussions. We thank Joshua Grant for proofreading the manuscript. We thank Ahmet Nihat Simsek for assisting with siibra software.

## Author’s contributions

AS - Conceptualization, Methodology, Software, Formal Analysis, Writing – Original Draft, Writing – Review & Editing; FS - Methodology, Writing – Review & Editing; LJE - Methodology, Software, Writing – Review & Editing; ALS - Software, Writing – Review & Editing; SH - Writing – Review & Editing; BM - Methodology, Writing – Review & Editing; KJP - Methodology, Software; KA - Conceptualization, Writing – Review & Editing; EK - Conceptualization, Methodology, Writing – Review & Editing; NW - Writing – Review & Editing; AV - Supervision, Writing – Review & Editing; VN - Conceptualization, Methodology, Supervision, Writing – Review & Editing

## Code availability

The code that was used for the study is available at https://github.com/astudenova/MEEG-cyto-myelo.

## Data availability

The LEMON dataset is publicly available at https://fcon_1000.projects.nitrc.org/indi/retro/MPI_LEMON.html.

The Cam-CAN dataset is publicly available at https://camcan-archive.mrc-cbu.cam.ac.uk/dataaccess/.

The BigBrain and Julich brain atlas can be explored at https://atlases.ebrains.eu/ or downloaded from BigBrainWarp toolbox https://bigbrainwarp.readthedocs.io/en/latest/ or siibra-python toolbox https://siibra-python.readthedocs.io/en/latest/.

The 7T dataset and Multimodal dataset cannot be made publicly available due to data protection. Aggregate data will be available at OSF upon publication.

