## Supplementary material for "Inter- and intra-individual variability in structure-function coupling in human brain": Inter- and intra-individual variability in structure-function coupling in human brain-38-48.pdf

### Supplementary information

Table S1. Short overview of the empirical studies revealing the relation between alpha rhythm power (amplitude) and layer-wise cortical microstructure

| Ref | Procedure | Species | Method | Brain dynamics | Type | Finding |
|---|---|---|---|---|---|---|
| Neymotin et al., 2020 | *in-silico* | - | computational model (HNN) | alpha power, alpha frequency | inter-layer | alpha rhythm shape and frequency resembled real data if pyramidal neurons were located in layers II, III, and V (which in the real brain do contain the majority of pyramidal neurons), and the network received both feedforward and feedback inputs |
| Traub et al., 2020 | *in-vitro* | rats | brain slices | alpha power | inter-layer | alpha rhythm power was correlated with bursting in layers IV and V |
| Bollimunta et al., 2008 | *in-vivo* | macaque monkeys | deep electrodes | alpha power | inter-layer | local pacemakers of the alpha rhythm were found both in the superficial and deep layers of visual areas |
| Haegens et al., 2015 | *in-vivo* | macaque monkeys | deep electrodes | alpha power | inter-layer | stronger generators of alpha rhythm were found in supragranular layers of primary auditory, visual, and somatosensory areas |
| Lopes da Silva et al., 1977 | *in-vivo* | dogs | deep electrodes | alpha power | inter-layer | alpha rhythm had higher power in layer V |
| Buffalo et al., 2011 | *in-vivo* | rhesus monkeys | deep electrodes | alpha power | inter-layer | attention affected alpha power in deep layers |
| Halgren et al., 2019 | *in-vivo* | human patients with epilepsy | ECoG | alpha power | inter-layer | alpha rhythm was found to have higher power primarily in supragranular layers |
| Goldman et al., 2002 | *in-vivo* | humans | combined fMRI-EEG | alpha power | intra-individual | the increase in alpha power over occipital sites was correlated with an increase in the signal from the thalamus |

| | | | | | | |
|---|---|---|---|---|---|---|
| Moosmann et al., 2003 | *in-vivo* | humans | combined fMRI-EEG | alpha power | intra-individual | positive correlation between occipital alpha power and fMRI signal from the thalamus, maximal at the left dorsomedial thalamic nucleus |
| Schreckenberger et al., 2004 | *in-vivo* | humans | combined PET-EEG | alpha power | intra-individual | alpha power across the cortex was reduced when glucose metabolism was reduced in the thalamus |
| Lindgren et al., 1999 | *in-vivo* | humans | combined PET-EEG | alpha power | intra-individual | alpha power was correlated negatively with metabolism in the thalamus |
| Kumral et al., 2022 | *in-vivo* | humans | EEG and structural MRI | alpha power | inter-individual | alpha power was shown to be increased with the decrease of thalamocortical connections caused by white matter degeneration |
| Shafiei et al., 2023 | *in-vivo* | humans | EEG and post-mortem brain | alpha power | intra-individual | features of the MEG signal, including alpha power, were correlated with the thickness of layer IV |
| Mahjoory et al., 2020 | *in-vivo* | humans | EEG and structural MRI | alpha frequency | intra-individual | alpha frequency was correlated with total cortical thickness |
| Sherman et al., 2016 | *in-silico*, *in-vivo* | humans | MEG, HNN | beta waveform | inter-layer | some parts of the beta cycle were induced by activity in more superficial layers, while others were induced by activity in deep layers |
| Mitchell et al., 1980; Alonso et al., 1987 | *in-vivo* | rats | deep electrodes | theta power | inter-layer | hippocampal theta was shown to have higher power in superficial layers |
| Artemenko, 1972 | *in-vivo* | rabbits | deep electrodes | theta power | inter-layer | theta power was higher in the middle layers of the hippocampus |
| Halgren et al., 2015; Halgren et al., 2018 | *in-vivo* | human patients with epilepsy | laminar microelectrode arrays | theta power | inter-layer | theta power was higher in the middle and superficial layers in different locations in the inferotemporal, perirhinal, prefrontal, and anterior cingulate cortices |

| | | | | | | |
|---|---|---|---|---|---|---|
| Halgren et al., 2021 | *in-vivo* | monkeys, mice, and human patients with epilepsy | laminar microelectrode arrays | 1/f slope | inter-layer | 1/f slope was steeper in the superficial layers in comparison to the deep layers |

Electrocorticography - ECoG, functional magnetic resonance imaging - fMRI, electroencephalography - EEG, positron emission tomography - PET, magnetic resonance imaging - fMRI, Human Neocortical Neurosolver - HNN

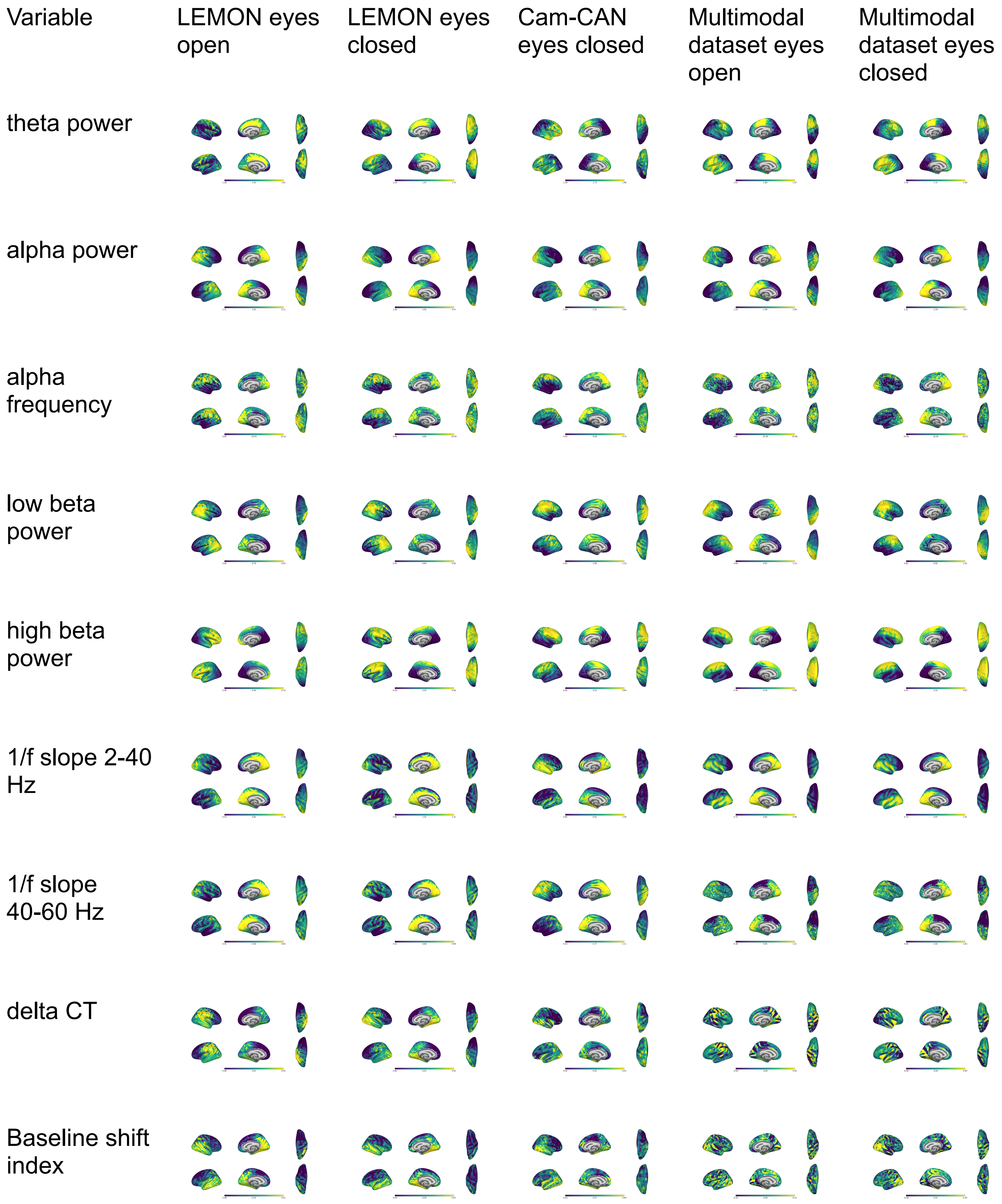


MPM superficial

MPM mid-cortical

MPM deep

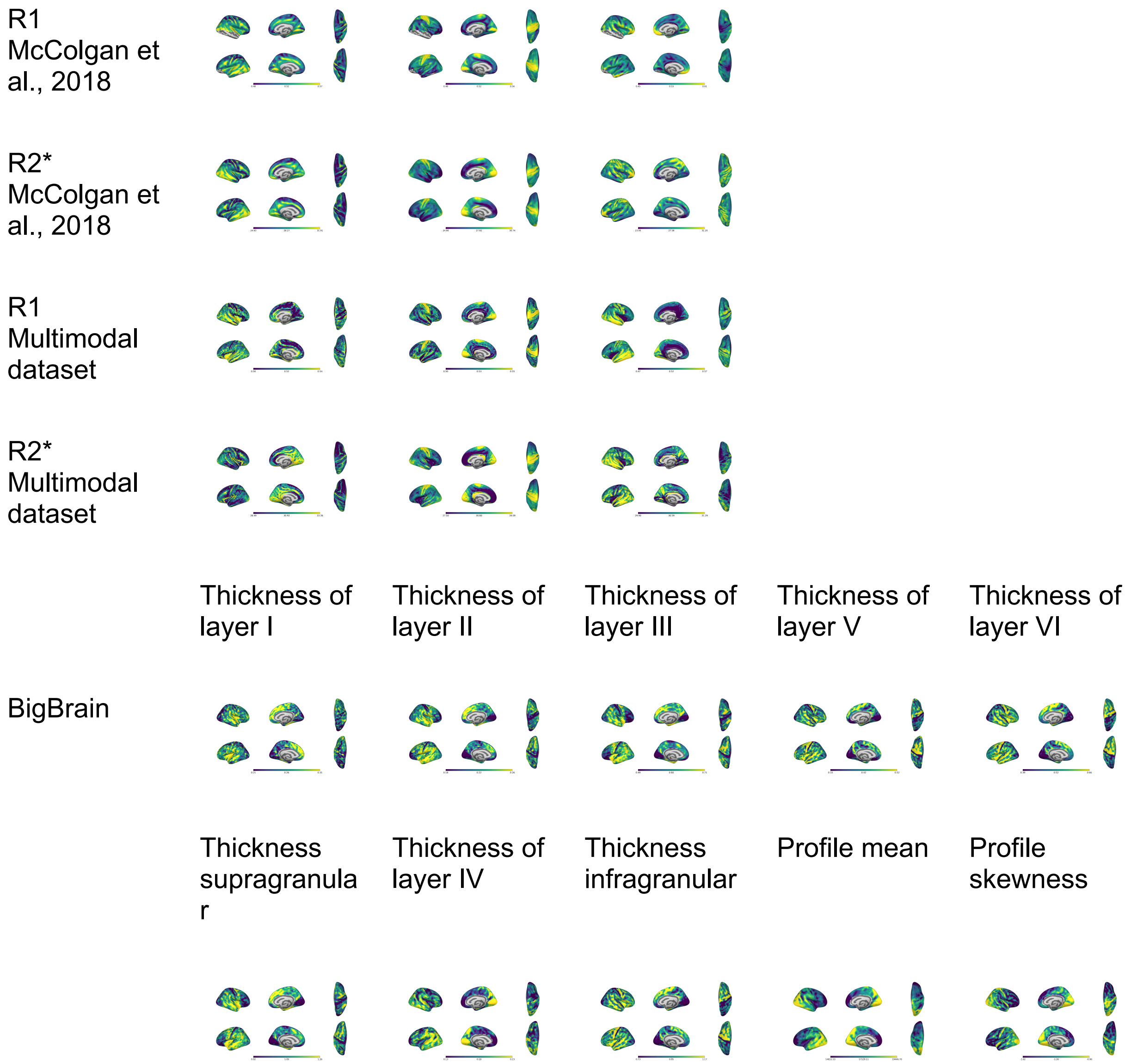


Figure S1. Distribution of MEG and EEG variables and correlation between two modalities, as well as cytoarchitectonic and myelin estimates. Some plots are duplicated from fig. 2.

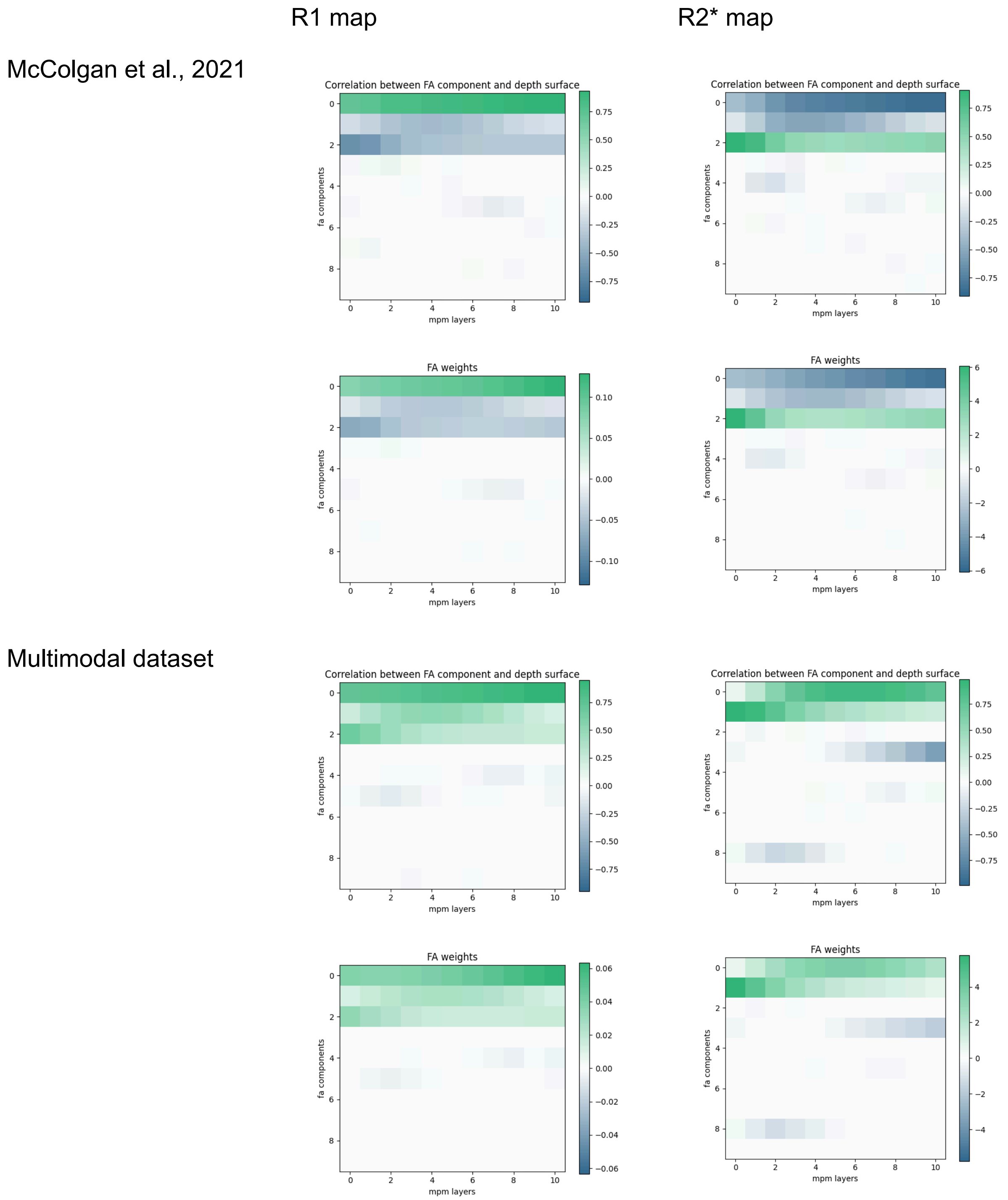


Figure S2. Factor analysis. The correlation between the obtained factors and the original depth surfaces. The 0th MPM layer is the pial surface, the 10th MPM layer is the gray to white matter border. We selected components that had a peak correlation of more than 0.3. Correlation direction is arbitrary. For the dataset from McColgan et al. (2021), the first three factors for both R1 and R2* were used. For the multimodal dataset, R1 first three factors were used, R2* first, second, and fourth factors were used.

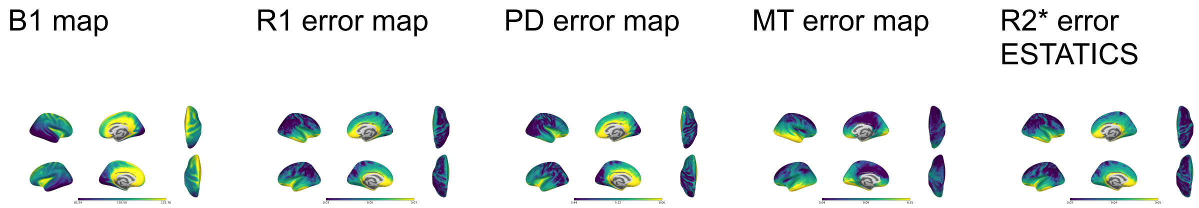


Figure S3. B1 field, and error maps from the Multimodal dataset. Error maps allow the estimation of variation of the noise from different factors, for instance, head coil configuration, different acquisition protocols, and artefacts such as head motion.

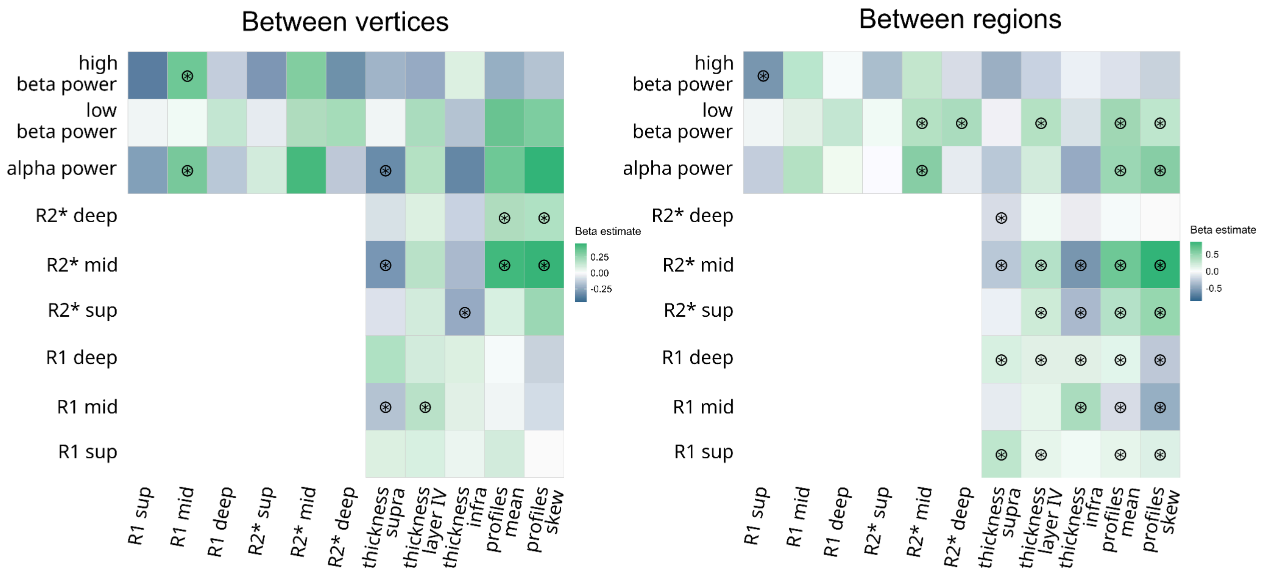


Figure S4. Exploratory analysis in the Multimodal dataset (same analysis as in Fig. 3C). The correlations are of similar direction with some differences. The absence of significance for some correlations may be explained by a smaller sample size (31 participants) and lower resolution of MRI data.

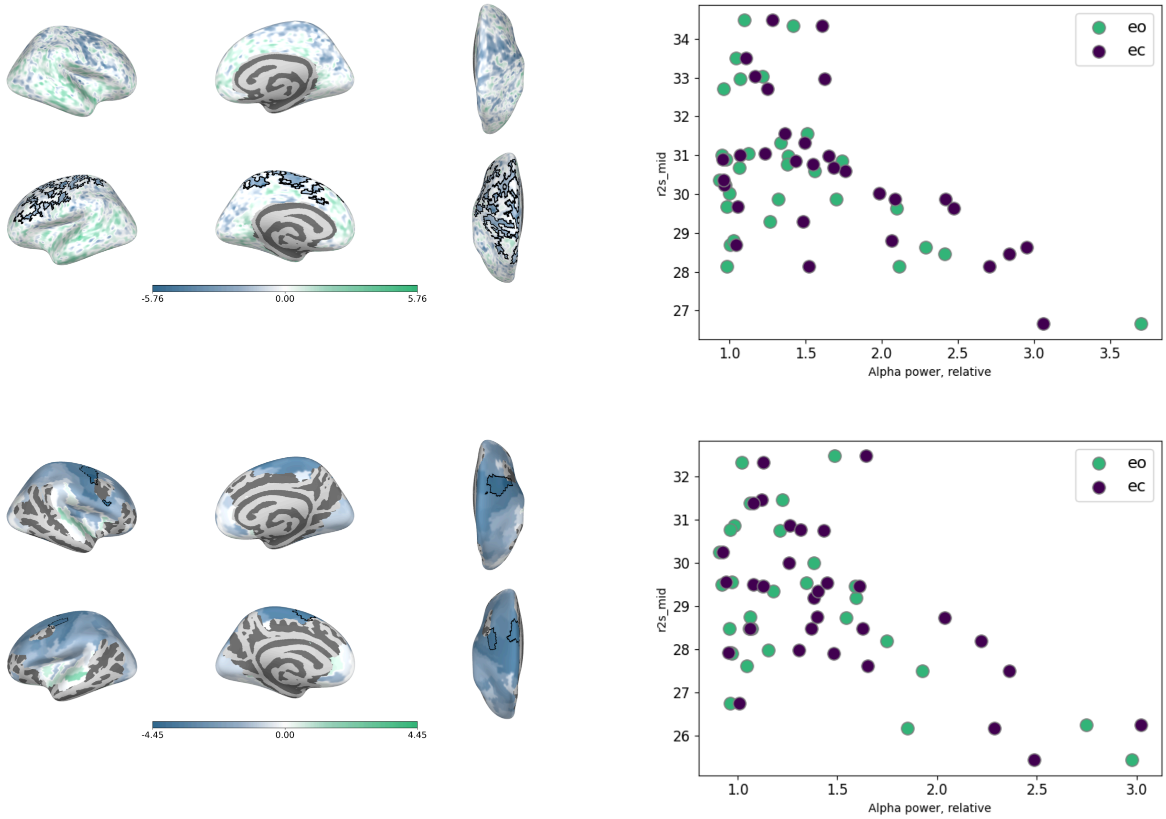


Figure S5. Between participants correlation between R2* and alpha power for the mid-cortical depth surface. Scatterplots are averaged for all the vertices (regions) that demonstrated significant correlation.

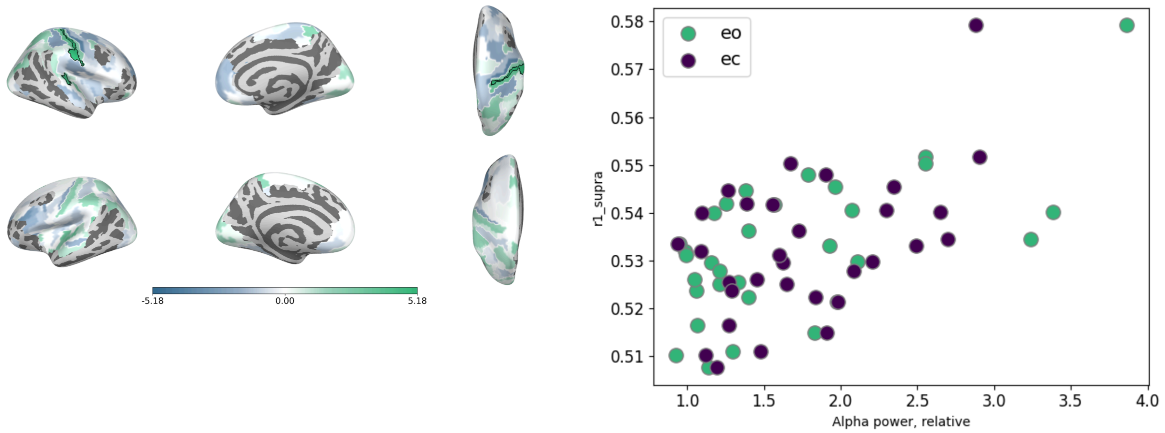


Figure S6. Significant association with R1 superficial and alpha power. Only found in analysis using parcellated data. Scatterplots are the averaged values within two significant regions.
